# Modulation of ribosomal subunit associations by eIF6 is critical for mitotic exit and cancer progression

**DOI:** 10.1101/2024.06.24.600220

**Authors:** Poonam Roshan, Aparna Biswas, Stella Anagnos, Riley Luebbers, Kavya Harish, Sinthyia Ahmed, Megan Li, Nicholas Nguyen, Gao Zhou, Frank Tedeschi, Vivian Hathuc, Zhenguo Lin, Zachary Hamilton, Sofia Origanti

**Affiliations:** Department of Biology, Saint Louis University, Saint Louis, MO, USA; Division of Urologic Surgery, Saint Louis University School of Medicine, Saint Louis, MO, USA; Center for RNA Science and Therapeutics, School of Medicine, Case Western Reserve University, Cleveland, OH, USA; Department of Pathology, Saint Louis University School of Medicine, Saint Louis, MO, USA

**Keywords:** eIF6, 60S, mitotic translation, vacant 80S, chromosome segregation, cancer invasion

## Abstract

Moderating the pool of active ribosomal subunits is critical for maintaining global translation rates. A factor crucial for modulating the 60S ribosomal subunits is eukaryotic translation initiation factor 6. Release of eIF6 from 60S is essential to permit 60S interactions with 40S. Here, using the N106S mutant of eIF6, we show that disrupting eIF6 interaction with 60S leads to an increase in vacant 80S. It further highlights a dichotomy in the anti-association activity of eIF6 that is distinct from its role in 60S biogenesis and shows that the nucleolar localization of eIF6 is not dependent on uL14-BCCIP interactions. Limiting active ribosomal pools markedly deregulates translation especially in mitosis and leads to chromosome segregation defects, mitotic exit delays and mitotic catastrophe. Ribo-Seq analysis of the eIF6-N106S mutant shows a significant downregulation in the translation efficiencies of mitotic factors and specifically transcripts with long 3′UTRs. eIF6-N106S mutation also limits cancer invasion, and this role is correlated with the overexpression of eIF6 only in high-grade invasive cancers suggesting that deregulation of eIF6 is probably not an early event in cancers. Thus, this study highlights the segregation of eIF6 functions and its role in moderating 80S availability for mitotic translation and cancer progression.

## Introduction

Assembly of the ribosomal subunits is a complex process that spans the nucleolus to cytoplasm and requires several conserved *trans*-acting factors^1–4^. Eukaryotic translation initiation factor-6 (eIF6) is one such essential factor that is critical for 60S biogenesis and for moderating inter-subunit interactions^5–10^. eIF6 functions as an anti-association factor that sterically inhibits 60S association with the 40S subunit through disruption of inter-subunit bridges^11–15^. Release of eIF6 from 60S is thus critical to permit 60S interactions with 40S to form the 80S ribosome^11–16^. Complete loss of eIF6 is lethal and results in a decrease in total 60S levels due to defects in pre-rRNA processing^7,10^. However, partial loss of eIF6 as seen in the heterozygous knockout mice does not affect normal growth or basal translation rates but leads to high levels of vacant 80S (devoid of mRNA) that impairs insulin-stimulated or growth factor-stimulated protein synthesis^7,17,18^. The high levels of vacant 80S are due to the spurious associations between 60S and 40S in the absence of sufficient cytoplasmic levels of eIF6. Thus, regulating the levels of eIF6 and its release from 60S at the temporal and spatial scale is important to modulate translation rates by establishing an equilibrium in the pools of active 60S and 80S that are available to engage in translation.

A deregulation of eIF6 activity is observed in inherited ribosomopathies such as the Shwachman-Diamond syndrome (SDS) and several cancers^12,19–24^. In SDS, release of eIF6 from 60S is hindered that limits the pool of freely available 60S^12,19,20,25–27^. This leads to a global decrease in translation rates^12,19,20,25–28^. Therefore, disrupting the interactions between eIF6 and 60S has been proposed as a therapeutic strategy to alleviate the disease phenotype especially the leukemic predisposition of SDS patients^19,26,29,30^. However, we are yet to completely understand the functional importance of targeting the direct interactions between eIF6 and 60S, especially in cancers.

eIF6 is overexpressed in many cancers including colon, ovarian, head and neck cancers and its enhanced expression is correlated with a poor prognosis^7,9,18,21,23,31,32^. Knocking down eIF6 in malignant pleural mesotheliomas and in Myc-induced lymphomas markedly impairs cell growth by inhibiting protein synthesis rates^7,9,18^. Heterozygous knockout of eIF6 markedly delays the transformation potential of oncogenic H-Ras12V mutant and oncogenic Myc expression^7,18^. Since partial loss of eIF6 delays tumorigenesis without markedly affecting normal growth and basal translation rates in normal cells, it presents eIF6 as a potent therapeutic target that is selective for cancers^7,18^. However, it remains unclear as to how high levels of eIF6 alter its interactions with 60S to sustain tumor growth and progression.

To better understand the crucial interactions between eIF6 and 60S, and their influence on cancer growth, we previously identified the key interface of interaction and showed that the terminal 8 residues in the C-tail of uL14 (RPL23) of 60S are essential for interactions with eIF6^30^. Mutation of several key residues in eIF6, especially N106S, were shown to disrupt interactions with RPL23 both *in vitro* and in cells^30^. Intriguingly, eIF6-N106S is also a predominant somatic mutation found in SDS patients^26,29^. Disrupting eIF6-60S interactions through the N106S mutation in SDS patients is considered to be beneficial for rescuing the growth and translation fitness of SDS disease^26,29^.

While we understand the importance of disrupting the eIF6 and 60S interaction interface for SDS phenotype, we are yet to understand its significance for cancers. To address the mechanism involved, we previously generated homozygous eIF6-N106S knock-in mutant cells and showed that disrupting the eIF6-60S interactions markedly slows the proliferation of colonic cancer cells with a mild increase in apoptosis^30^. However, the mechanistic effect of disrupting the direct interaction between eIF6 and 60S on cell proliferation and translation profiles in cancers remains unclear. In addition, though it has been known for decades that eIF6 is important for cell proliferation, the mechanism of how eIF6 regulates cell cycle progression remains unknown.

Here, we show that disrupting eIF6 and 60S interactions through the N106S mutation leads to an increase in vacant 80S ribosomes and a decrease in global translation rates. This ribosomal stress triggers the p53-checkpoint pathway independent of DNA damage. The N106S mutation, however, does not affect total 60S levels suggesting that eIF6 role in nucleolar 60S biogenesis is distinct from its anti-association activity. We also show that the nucleolar localization of eIF6 is preserved in mutants that is sufficient to maintain total 60S levels. Contrary to a previous model, we show that the nucleolar localization of eIF6 is not dependent on its direct interactions with the BCCIP chaperone. Altering 80S monosome availability is rate limiting for sustaining global translation rates especially in mitosis but not in other phases of cell cycle. The deregulated mitotic translation rates in the eIF6-N106S mutant cells hinder chromosome alignment and chromosome segregation that delays cytokinesis and induces mitotic catastrophe. Ribo-seq analysis shows a significant downregulation in the translation efficiency (TE) of transcripts associated with centromeres, chromosome localization and for transcripts with long 3’UTRs. Ribo-Seq analysis also shows altered TEs for transcripts associated with cancer invasion and cytoskeleton, and we observe that the N106S mutation markedly limits cancer growth and invasion *in vitro* and in xenografts *in vivo.* The marked effect of eIF6 on cancer invasion is further corroborated by overexpression of eIF6 only in high-grade invasive bladder cancers but not in low-grade cancers or benign tissues derived from patients suggesting that deregulation of eIF6 is not an early event in cancers.

## Results

### Activation of the p53-mediated cell cycle checkpoint response in the eIF6-N106S mutant cells

Our previous study identified the importance of the C-tail of uL14 of 60S in mediating interactions with the N106 residue of eIF6 (Fig. 1A)^30^. To better understand the cellular importance of interactions, we had previously generated the homozygous *eIF6-N106S* knock-in mutation in HCT116 cells using CRISPR gene editing^30^. Expression of the eIF6-N106S mutant is similar to WT with only a mild decrease in levels (Fig. 1B), as shown previously^30^. We previously showed that the N106S mutation markedly decreases the proliferation rate of the colonic cancer cells with only a mild increase in apoptosis as determined by the cleaved caspase-3 assay^30^. These results provided the first proof-of-concept for therapeutically disrupting eIF6 and uL14 interactions to inhibit cancer cell proliferation. However, the mechanism of how cell proliferation is disrupted in these mutant cells and how eIF6 affects cell cycle progression remains unknown.

**Figure 1.**
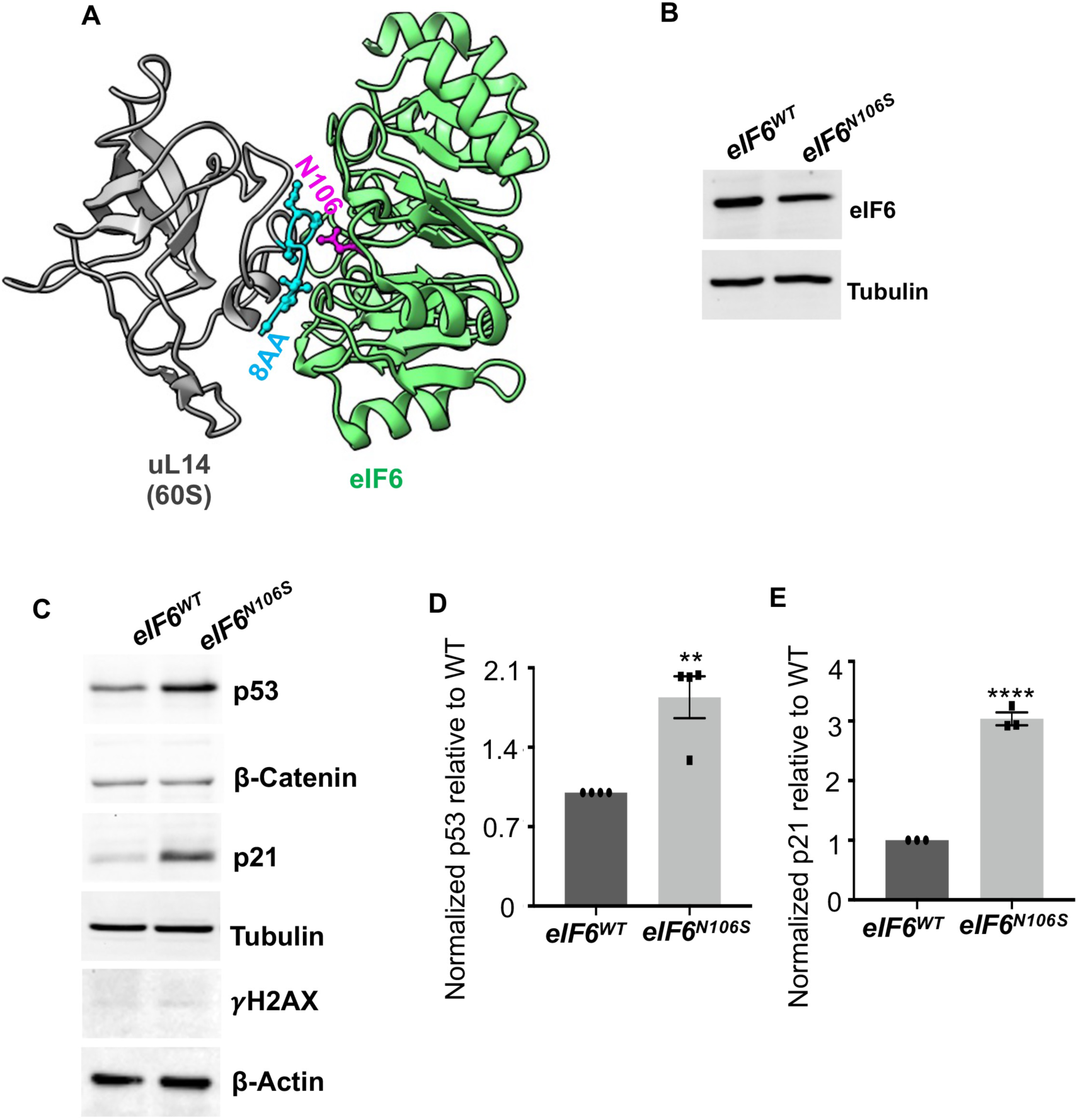
Disruption of eIF6 and uL14 interactions through N106S mutation induces p53 checkpoint response. A) Cryo-EM structure of human eIF6 (green) interacting with uL14 (RPL23) (gray) of pre-60S ribosomal subunit (PBD: 6LU8). The key residues of interaction in eIF6 (N106) (pink) and the terminal 8 residues (8AA) in uL14 (blue) are highlighted. B) Representative western blot of HCT116 cells with homozygous knock-in of eIF6-N106S mutation in comparison to isogenic WT controls. Blots probed with anti-eIF6 antibody (Santa Cruz Biotechnology) and anti-β-Tubulin antibody as loading control C) Representative western blots was probed with anti-p53, anti-p21 and anti-𝛾H2AX antibodies. β-Tubulin, β-Catenin and β-Actin used as loading controls. D and E) Blots represented in C. were quantitated and normalized to loading control. Plots represent standard error of the mean of four (D) and three (E) independent experiments. Significant differences in p53 (*p*=0.0039) (D) and p21 levels (*p*=0.000047) (E) relative to WT were determined by an unpaired two-tailed *t* test.

To address the effects on cell cycle progression, we assessed activation of the p53-mediated cell cycle checkpoint response in eIF6-N106S mutant cells. We found that the levels of both p53 and its transcriptional effector: p21 are induced in the N106S mutant (Fig. 1C, D and E). p53 can be induced by ribosomal stress response that is independent of its activation by the DNA damage response^33^. To determine if p53 is induced in the mutants due to a DNA damage response, we probed for changes in **γ**H2AX where H2AX, a histone variant, is phosphorylated at Ser139 in response to DNA damage, especially in response to DNA double strand breaks^34^. However, we did not observe any upregulation of **γ**H2AX (Fig. 1C). This suggests that activation of the p53-checkpoint response is primarily attributed to ribosomal stress.

### Increased vacant 80S levels and translation repression upon disruption of eIF6-60S interaction

To better understand the ribosomal stress, we assessed the polysome profiles of the eIF6-N106S mutant cells. The mutants showed high levels of vacant 80S and a decrease in the polysome peak suggesting that global protein synthesis is markedly reduced in the mutant (Fig. 2A, B, C and D). The overall translation rates were also decreased in the mutant as determined by the click-chemistry based assay using methionine analog (L-Azidohomoalanine; AHA) (Fig. 2E). To determine the affinity of eIF6 mutant for 60S, we assessed the relative distribution of eIF6 in the non-ribosomal fractions compared to the 60S fractions (Fig. 2B). The corresponding input levels of eIF6-WT and eIF6-N106S are also shown (Fig. 2C). As expected, eIF6-WT bound to 60S however, the eIF6-N106S mutant was more abundant in the non-ribosomal fraction than the 60S fraction (Fig. 2B). Since we previously showed that the N106S mutant interacts poorly with uL14 (RPL23) of the 60S subunit^30^, these results indicate that disruption of uL14 binding is sufficient to hinder interactions with 60S. This also suggests that decreased binding of eIF6 to 60S leads to an increase in vacant 80S peak due to spurious associations between 60S and 40S in the absence of anti-association activity of eIF6. Also, we observed only a mild decrease in 40S levels and 60S levels in the mutant compared to WT suggesting that the effect of N106S mutation primarily affects the anti-association activity of eIF6 rather than eIF6 role in 60S biogenesis (Fig. S1A).

**Figure 2.**
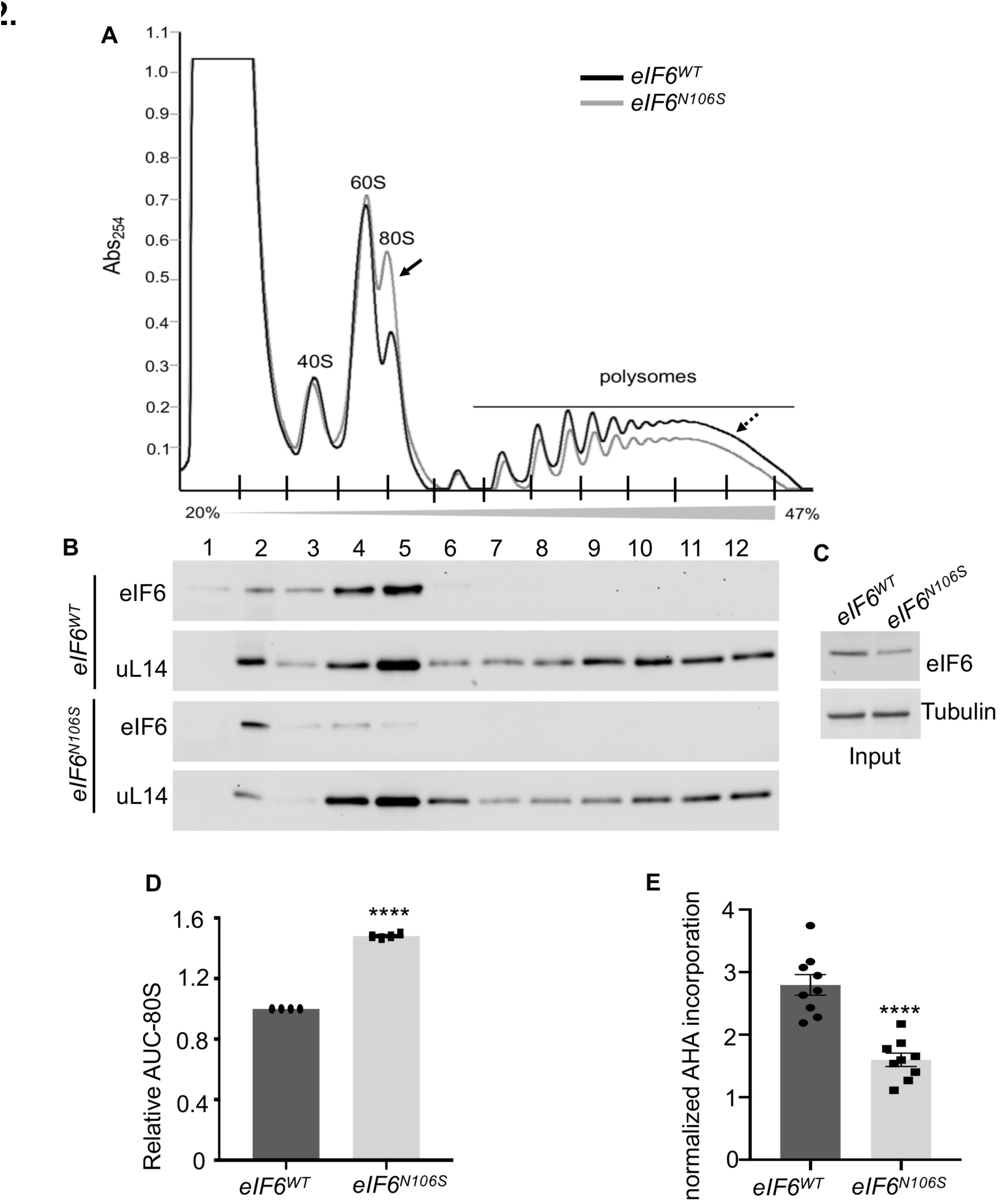
Increased empty 80S levels and reduced rate of global translation due to weakened binding of eIF6-N106S mutant to the 60S subunit. A) Representative polysome profiles of eIF6-N106S (gray) relative to eIF6-WT (black) HCT116 cells assayed by sucrose gradient centrifugation with a gradient of 20% to 47% sucrose. Arrows indicate differences in 80S peak (solid arrow) and polysome peak (dashed arrow). B) Representative western blots of proteins extracted from the indicated sucrose-gradient fractions. Blots represent three independent experiments. Blots were probed with anti-eIF6 and anti-uL14 antibodies. C) Western blot represents eIF6 and β-Tubulin (loading control) levels in the input for polysome profile assay. D) Plot shows the relative area under the curve (AUC) measurements for 80S levels in eIF6-WT and eIF6-N106S mutant. Plot shows standard error of the mean of four independent experiments with a significant difference between eIF6-WT and eIF6-N106S (p=0.0000000003) determined by an unpaired two-tailed *t* test. E) Plot shows normalized AHA incorporation rates assayed in triplicate per experiment and depicted as standard error of the mean of three independent experiments. Significant difference between eIF6-WT and eIF6-N106S (p=0.00001) determined using an unpaired two-tailed *t* test.

### Nucleolar localization of eIF6 is not dependent on BCCIP or uL14

eIF6 localization to distinct nucleoli and cytoplasmic compartments are attributed to its distinct roles in 60S biogenesis and anti-association activity^10,18^. We, therefore, probed the subcellular distribution of eIF6. We observed a ∼15% increase in the cytoplasmic localization of eIF6 and a corresponding decrease in the nuclear fraction (Fig. 3A, B and C). This was also observed in our immunofluorescence studies where endogenous eIF6 showed a decrease in overall nuclear staining of eIF6 in the mutant cells as shown by increased visibility of the nuclear (DAPI) stain (Fig. 3D). We also observed mild changes in overall cell morphology and cell to cell contacts in the mutant that was also observed by brightfield imaging and α-Tubulin staining (Fig. S1B). Remarkably, the nucleolar staining of eIF6 was intense suggesting that eIF6 localization to the nucleolus was mostly preserved in the mutant cells (Fig. 3D). Previous studies have shown that depleting the nucleolar pool of eIF6 disrupts its function in pre-rRNA processing and 60S biogenesis^10^. However, since eIF6-N106S mutant localizes to the nucleolus and the 60S levels are maintained in the mutant cells, it further emphasizes that the N106S mutation does not significantly affect 60S biogenesis. This also suggests that eIF6 interaction with uL14 is not critical for its nucleolar localization. A previous study indicated that nucleolar localization of eIF6 is dependent on the direct interaction of eIF6 with the BCCIP chaperone, specifically with the β-isoform^35^. We therefore immunoprecipitated endogenous eIF6 and probed for its interaction with BCCIP (Fig. 3E). eIF6-N106S mutant was not found to interact with BCCIP unlike the WT (Fig. 3E). Similar results were obtained by immunoprecipitating endogenous BCCIPβ and probing for its interaction with eIF6 (Fig. 3F). These results indicate that eIF6 interaction with BCCIP or uL14 is not critical for its nucleolar recruitment or retention.

**Figure 3.**
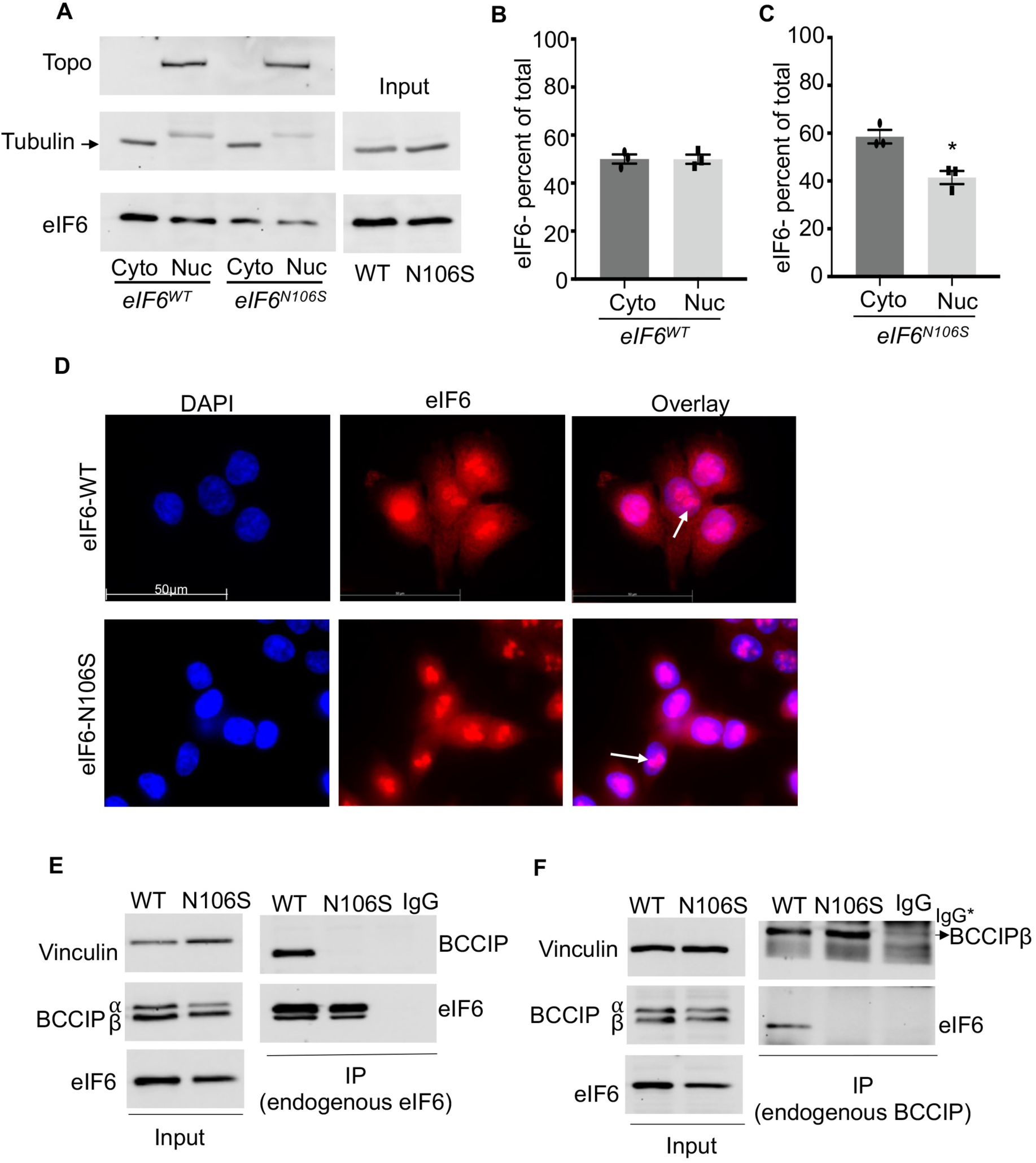
Nucleolar localization of eIF6 is not dependent on its interaction with BCCIP or uL14. A) Western blot represents the nuclear (nuc) and cytoplasmic (cyto) distribution of eIF6. Blots were probed with anti-eIF6 (Santa Cruz Biotechnology), anti-Topoisomerase II-β (nuclear marker) and anti-β-Tubulin (cytoplasmic marker) antibodies. Adjacent western blots represent the total input levels of eIF6 and β-Tubulin (loading control). B and C) Blots shown in A. were quantitated and the percent distribution of eIF6 in nuclear and cytoplasmic fractions were plotted for eIF6-WT (B) and eIF6-N106S mutant (C). Plots indicate the standard error of the mean of three independent experiments. D) IF images of asynchronous cells show localization of eIF6 to nucleoli (white arrows) in both WT and mutant cells and decreased nuclear localization in mutant. Fixed cells were stained with DAPI (nuclear marker) or incubated with anti-eIF6 antibody. E) Representative western blot shows immunoprecipitation of endogenous eIF6 or anti-mouse IgG control and its interaction with endogenous BCCIP. The corresponding input is shown. Vinculin used as loading control. F) Representative western blot shows immunoprecipitation of endogenous BCCIPβ or anti-rabbit IgG control and its interaction with endogenous eIF6. The corresponding input is shown. Vinculin used as loading control. (*Heavy-chain IgG migrates close to BCCIPβ in the gel.)

### eIF6-N106S mutation deregulates mitotic translation and mitotic progression

Given the marked inhibition of global translation, we wanted to determine if the effects on translation could be attributed to deregulation of other translation initiation factors. No changes were observed in the levels of 4EBP1, 4EBP2, eIF2A, eIF4G1, eIF4G2, eIF4E, eIF2α, and Ser-209 phosphorylation of eIF4E (Fig. 4A, B, C, and Fig. S1C). The total 4EBP1 blot also showed hyperphosphorylated forms, however, no changes were observed in the hyperphosphorylated forms of 4EBP1. We also tested for changes in induction of Ser51-phosphorylation of eIF2α in response to oxidative stress. No changes were observed in the levels of Ser51-phosphorylation of eIF2α in the mutant cells compared to WT (Fig. 4C). Also, no compensatory changes in the levels of eIF6 release factors: SBDS and EFL1-GTPase were observed (Fig. 4D). These results indicate that the translation effects are primarily attributed to the deregulation of eIF6 and 80S ribosome availability rather than changes in initiation rates through modulation of eIFs.

**Figure 4.**
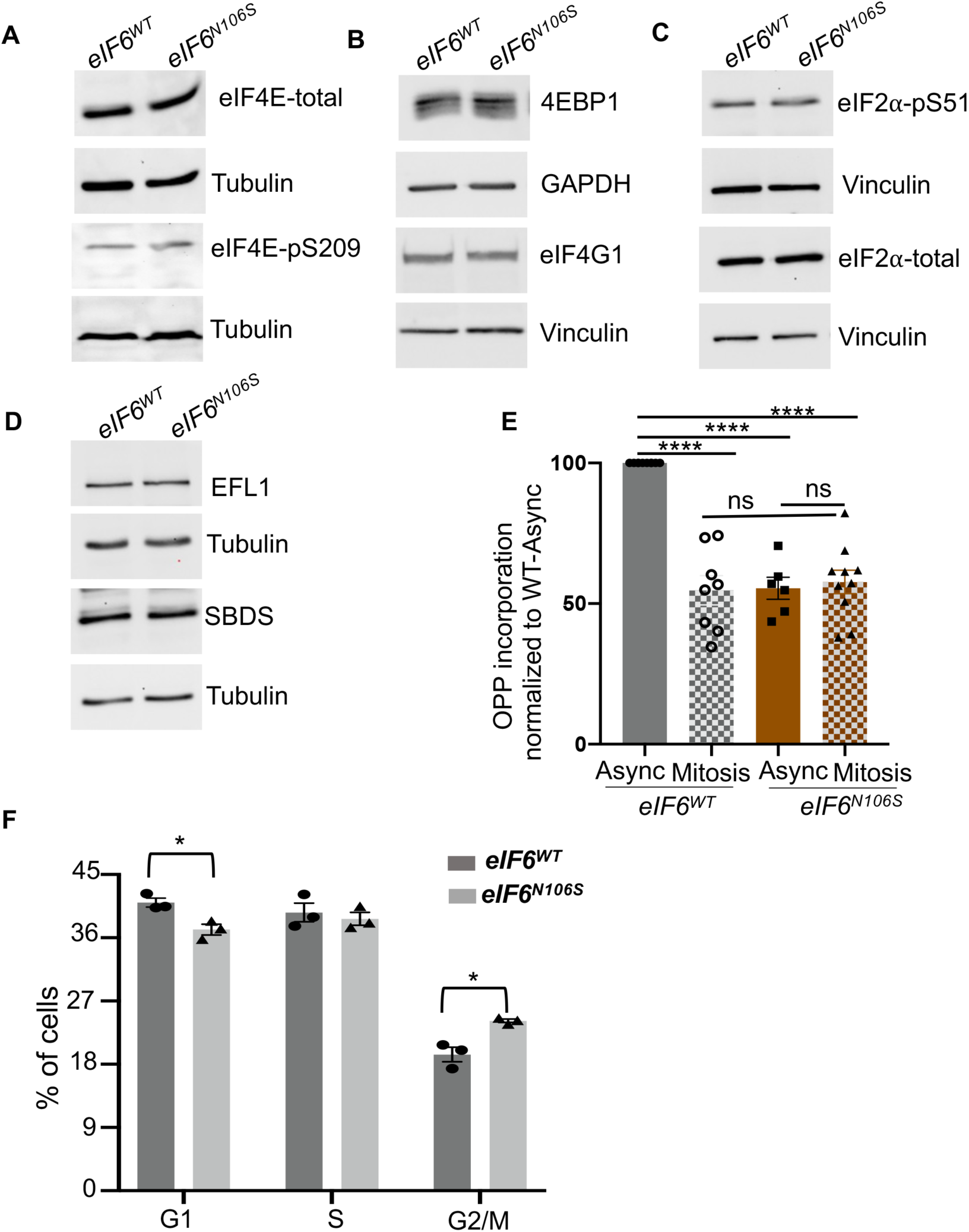
Deregulation of mitotic translation but unaltered regulation of other eIFs in eIF6-N106S mutant. A, B, C and D) Representative western blots of eIF6-WT and eIF6-N106S probed with the indicated antibodies. Blots were also probed with anti-Vinculin, anti-β-Tubulin or anti-GAPDH antibodies as loading controls. Serine-51 phosphorylation of eIF2⍺ was induced by oxidative stress with 4mM H_2_O_2_ for 1 hour. E) Global translation rates in asynchronous cells (Async) or mitotic cells synchronized with nocodazole were measured using the OPP incorporation assay. Plots represent the standard error of the mean of OPP incorporation rates of three independent experiments and normalized to the eIF6-WT asynchronous rates. Significant differences were found between WT-async and WT-mitosis (*p*=0.000000002), WT-async and N106S-async (*p*=0.000000021), and WT-async and N106S-mitosis (*p*=0.000000003) and no-significant (ns) differences were observed for N106S-async and N106S-mitosis (*p*=0.97) and WT-mitosis and N106S-mitosis (*p*=0.91) as determined by a one-way ANOVA and Sidak’s multiple comparisons test. F) Plot shows cell cycle profile analysis determined by propidium iodide staining and flow cytometry. Data depict the standard error of the mean of three independent experiments. Significant differences between eIF6-WT and eIF6-N106S observed for G1 phase (*p*=0.017) and G2/M phase (*p*=0.0105) of cell cycle as determined by an unpaired two-tailed *t* test.

Increased levels of vacant 80S are observed under physiological states such as mitosis and hibernation, or in response to stress induced by conditions such as nutrient deprivation^36–43^. The increased vacant 80S levels in mitosis are associated with a marked repression in global translation rates along with a decrease in translation rates of specific mRNAs^39–43^. Given the increased vacant 80S levels in the mutant cells, we wanted to determine the effect of eIF6 mutation on mitotic translation rates. We observed that translation rates in mitosis were reduced by 50% in the WT cells in comparison to asynchronous cells similar to previous reports (Fig. 4E)^39–43^. However, the translation rates that were already decreased in the asynchronous eIF6-N106S mutant cells, did not show any further decrease in mitosis (Fig. 4E). Repressing global translation is crucial to progress through mitosis^39–43^. However, if mitotic translation is deregulated, then it can markedly affect cytokinesis and mitotic exit as shown in cells lacking 14-3-3σ^40^. We therefore determined if mitotic progression was affected in the mutant cells. Analysis of cell cycle phases using flow cytometry showed that there is a mild but significant increase in the percentage of cells in G2/M phase of cell cycle with a corresponding decrease in the G1 population (Fig. 4F). This suggests that some cells are delayed in the G2/M phase of cell cycle that eventually delays entry into G1. Further analysis by immunofluorescence staining showed that the delay was predominantly in the mitotic phase of cell cycle (Fig. 5 and Fig. S2). There was a significant increase in the number of binucleate cells (Fig. 5A, B and S2A). There was also a significant increase in the number of cells exhibiting mitotic catastrophe as shown by presence of abnormally large cells with multinulcei, giant nuclei, micronuclei and vacuoles (Fig. 5C and D). The 25% to 30% increase in mitotic catastrophe is consistent with the extent of growth delay in the mutant.

**Figure 5.**
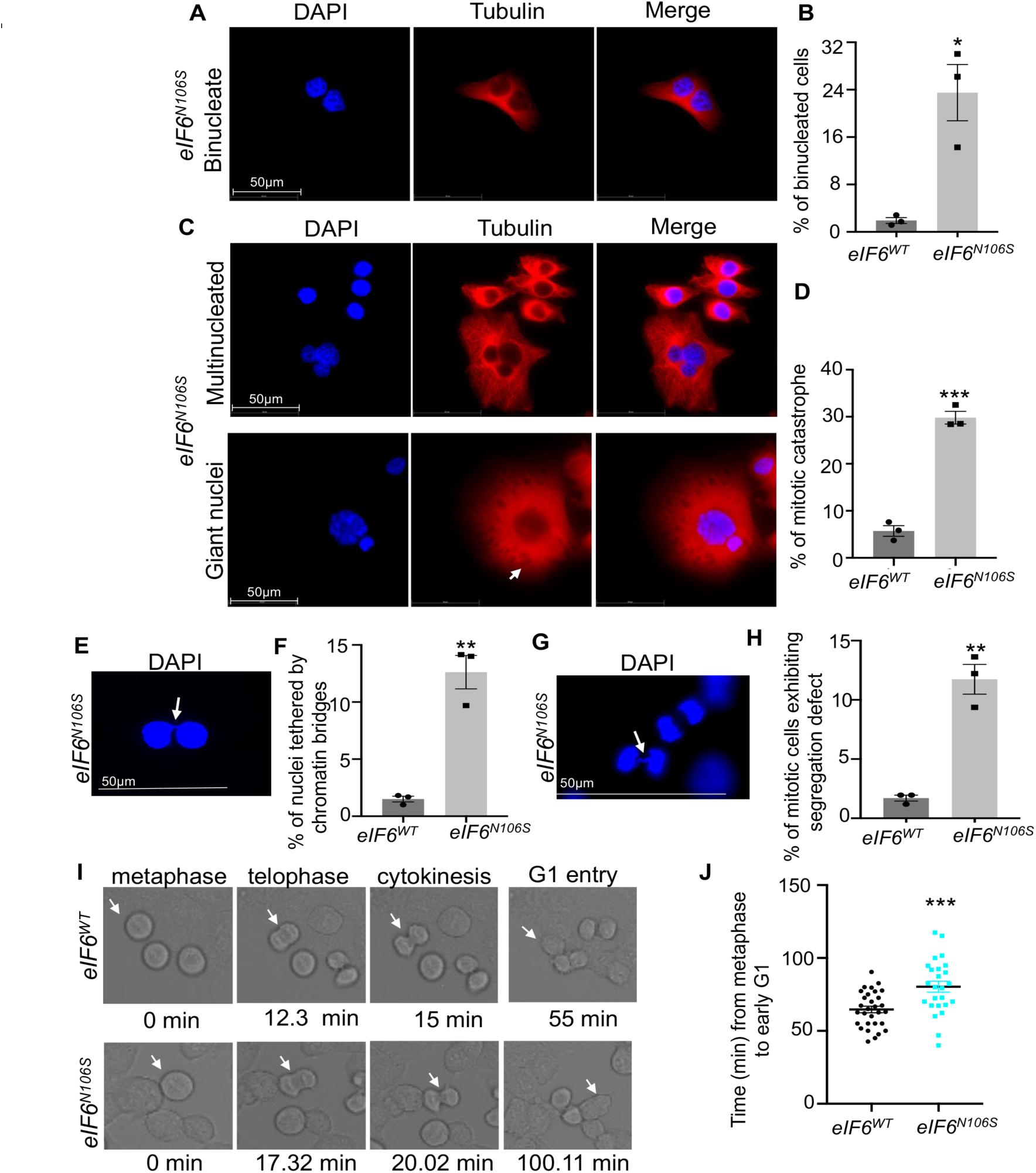
eIF6-N106S mutation leads to errors in chromosome segregation, mitotic exit delay and mitotic catastrophe. A and B) Representative immunofluorescence (IF) images show increased presence of binucleated cells for the eIF6-N106S mutant. Fixed asynchronous cells were stained with DAPI (nuclear stain) or treated with anti-Tubulin antibody. Plot (B) shows the standard error of the mean of binucleated cells counted from three independent experiments. A total of ∼900 cells were counted for each genotype for all three experiments combined. Statistical significance (*p*=0.0106) determined using an unpaired two tailed *t* test. C and D) Representative IF images show increased mitotic catastrophe with presence of cells with multinuclei, giant nuclei, micronuclei and vacuoles (white arrow). Plot (D) shows the standard error of the mean of giant and multinucleated cells counted from three independent experiments. Statistical significance (*p*=0.0002) determined using an unpaired two tailed *t* test. E and F) Representative IF image shows a binucleate cell where nuclei are tethered by a chromatin bridge (white arrow). Plot (F) shows the standard error of the mean of nuclei tethered by chromatin bridges during cytokinesis from a total of ∼400 asynchronous cells from three independent experiments and *p*=0.0017 as determined by an unpaired two-tailed *t* test. G and H) Representative IF images show increased presence of anaphase bridges in anaphase cells. Plot (H) shows the standard error of the mean of mitotic cells exhibiting segregation defects (anaphase bridges and/or lagging chromosomes) from three independent experiments. Statistical significance (*p*=0.0014) determined using an unpaired two tailed *t* test. I and J) Confocal time-lapse brightfield images show snapshots of cells (white arrows) progressing through metaphase to early G1. Cells were synchronized in G2 phase with RO-3306 (Cdk1 inhibitor) for 20 hours and then released into mitosis for imaging. The timing of progression from metaphase to G1-entry was measured and plotted. A total of 25 (eIF6-N106S) and 30 (eIF6-WT) cells undergoing mitosis were imaged. Statistical significance (*p*=0.0005) determined using an unpaired two tailed *t* test.

Intriguingly, most of the nuclei in cells undergoing cytokinesis were tethered by chromatin bridges that were indicative of chromosome segregation defects in the mutant cells (Fig. 5E, F and S2B). The eIF6-N106S mutant also showed a significant increase in anaphase bridges and lagging chromosomes in mitotic cells that further indicated chromosome segregation errors (Fig. 5G and H). However, cells synchronized in mitosis did not show any changes in the levels of Serine10-phosphorylation of Histone H3, a mitotic marker, indicating that entry into mitosis was not affected in the mutant cells (Fig. S2C). Also, no overt changes in the bipolarity of the spindles or spindle organization were observed (Fig. S2D). A second clone of eIF6-N106S mutant also showed similar delays in cell proliferation, mitotic catastrophe and altered cell morphology (Fig. S2E and F) suggesting that the effect of eIF6-N106S mutation on mitotic progression is not due to clonal variation. To further assess delays in mitotic progression, we synchronized cells in G2 phase and released them into mitosis followed by time-lapse live cell imaging (Fig. 5I, J and S3). Time-lapse imaging showed that while there is a minor delay in mitotic progression from metaphase to telophase, major delays were observed in cells progressing from cytokinesis to G1 in the eIF6-N106S mutant cells (Fig. 5I, J and S3). These results show that the deregulation of mitotic translation due to eIF6-N106S mutation causes defects in chromosome segregation that results in delays in cytokinesis and eventual cell death by mitotic catastrophe.

### Ribo-Seq analysis uncovers altered translational efficiencies of genes critical for mitosis in eIF6-N106S mutant

To better understand the translational deregulation that leads to mitotic defects, we performed ribosome profiling and RNA-Seq to measure the translation efficiencies of individual mRNAs as described before^44,45^. For the methodology of ribosome profiling, short fragments of mRNA bound by ribosomes and therefore, protected by a ribosome are released by digestion of the mRNAs with micrococcal nuclease^44,46^. This generates ribosome protected footprints (RPF) that determines the average number of ribosomes per mRNA protected fragments that are sequenced by the Illumina NovaSeq 6000. The mRNA levels are also quantified by RNA-Seq^44,46^. Translation efficiencies of individual mRNAs are then determined as the ratio of RPF/mRNA abundance. Using this methodology, we determined the global changes in translation efficiencies (ΔTE) of transcripts in the mutant compared to WT (Fig. 6 and Fig. S4, S5 and S6, Table S1). We focused on genes whose translation efficiencies (ΔTE) were downregulated or upregulated as 1.5-fold change (1.5FC) or more in mutant relative to WT (Fig. 6A and S6A). A global analysis of significantly downregulated ΔTEs showed that the changes in TEs were associated with a mild increase in total mRNA levels and a corresponding decrease in RPF abundance (Fig. 6B). A similar reciprocal regulation of significantly upregulated ΔTEs showed a mild decrease in total mRNA levels with a corresponding increase in RPF abundance (Fig. 6C). This suggests that some transcriptional rewiring triggered by changes in translational rates contributes to the altered TEs in the mutant cells. This was also corroborated by the GO-enrichment (Gene Ontology/KEGG) analysis which showed that genes associated with transcription factor activity were significantly downregulated by the N106S mutation (Fig. 6D).

**Figure 6.**
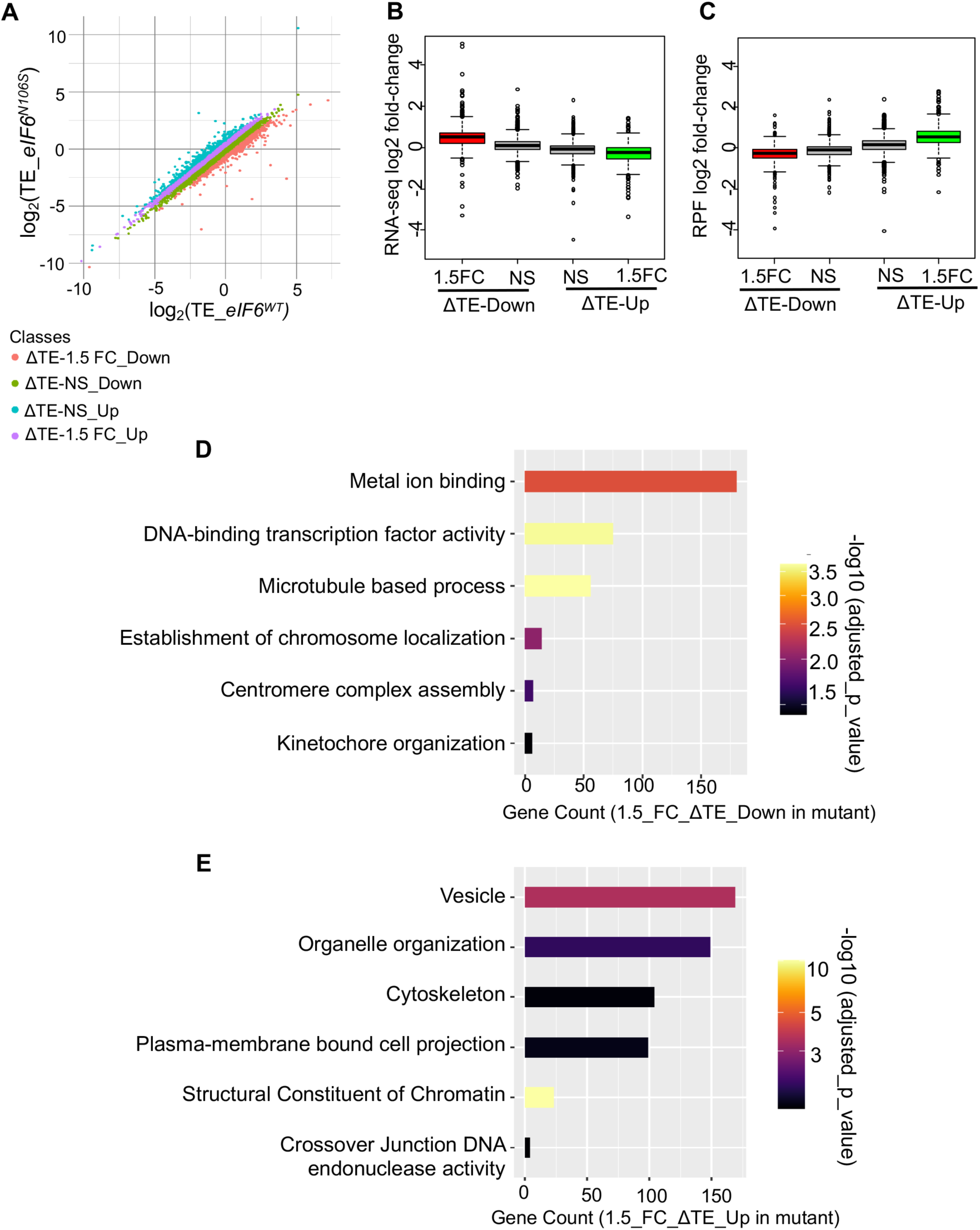
Ribosome profile analysis shows altered TEs of transcripts associated with mitotic kinetochore organization, chromosome localization and cytoskeleton in the eIF6-N106S mutant. A) Scatter plot shows the distribution of transcripts with a significant downregulation or upregulation in translation efficiencies (TEs) of 1.5-fold or more in mutant compared to WT (ΔTE) and non-significant ΔTEs. ΔTE calculations are indicated in supplementary. B and C) Box plot shows changes in levels of mRNA (B) and Ribosome Protected Footprints (RPF) (C) for transcripts with a significant 1.5-fold or more upregulation (green boxes) or downregulation (red boxes) in TEs in mutant compared to WT. Plots also depict non-significant ΔTEs (gray boxes). ANOVA test shows significant differences among the four classes (*p* < 2e-16) and significant differences between each group (*p* < 2e-16) according to Tukey’s HSD post hoc test. D and E) GO/KEGG-enrichment analysis was carried out on transcripts that were downregulated (D) or upregulated (E) by 1.5-fold or more in the mutant compared to WT. *p*-values were determined using Fisher’s one-tailed *t t*est performed by g:profiler. Read lengths for ribosome profiling and sample size are indicated in the supplementary.

GO-enrichment analysis revealed a marked downregulation of genes associated with mitotic kinetochores and centromeric assembly and chromosome localization in the mutant cells (Fig. 6D). Defects in kinetochore organization can lead to misalignment of chromosomes. Staining with anti-centromere antibody further showed that the chromosomes were often misaligned in the mitotic cells of the eIF6-N106S mutant (Fig. S7). This suggests that a translational deregulation of the factors critical for centromere and kinetochore organization are likely to hinder efficient chromosome alignment in mitosis and lead to the segregation defects as seen in the eIF6-N106S mutant cells. A marked increase in TEs of many the nucleosome components including histone variants were also observed suggesting that chromatin alterations could further contribute to the defects in mitotic progression and altered transcriptional activity (Fig. 6E).

Since the vacant 80S ribosomes are rate-limiting in the eIF6-N106S mutant, we wanted to determine if the translation efficiencies of longer transcripts were down regulated due to insufficient levels of active ribosomes. For this analysis, we focused only on transcripts that showed significant changes in TEs without any significant changes in mRNA levels. Intriguingly, the TEs of transcripts with long 3’UTRs and lower GC content were significantly downregulated in the mutant (Fig. 7A and B). Altered ribosomal availability has been shown to modulate 3′UTR ribosomal occupancy and recycling in certain mutants of ribosome rescue factors such as Dom34^47,48^. Given that we have promoted re-association of ribosomal subunits by targeting eIF6, it is possible that ribosomal occupancy on longer 3′UTRs and ribosome recycling have been altered as a consequence and hinder translation. Future studies will determine the effects of eIF6 mutation on 3′UTR ribosomal occupancy and recycling.

**Figure 7.**
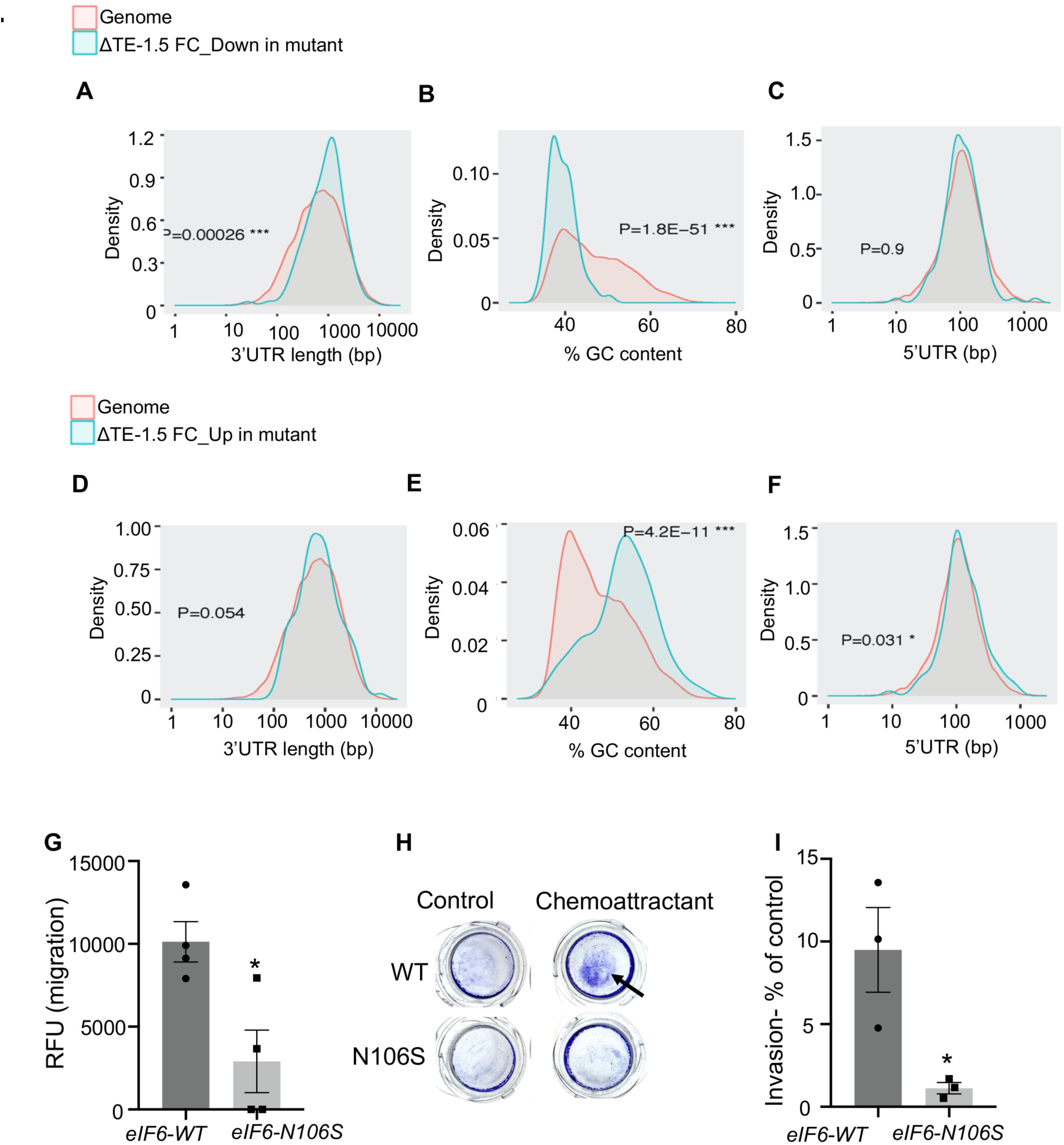
Downregulation of TEs for transcripts with long 3’UTRs and lower GC content and upregulation of TEs for transcripts with long 5’UTRs and high GC content. A, B and C) Density plots show the density of transcripts that are signicantly downregulated by 1.5-fold or more in mutant compared to WT and relative to the genome. Comparisons of density of transcripts based on 3’UTR length (A), percent GC content (B) and 5’UTR length are shown. D, E and F) Density plots show the density of transcripts that are significantly upregulated by 1.5-fold or more in mutant compared to WT and relative to the genome. Density comparisons of transcripts based on 3’UTR length (D), percent GC content (E) and 5’UTR length (F) are shown. *p*-values for all density plots were calculated using Student’s t-tests using ShinyGO 0.80 analysis. G) Plot depicts cell migration measured as relative fluorescence units (RFU) determined using a transwell migration assay with 10% serum as chemoattractant. Plot includes the standard error of the mean from four independent experiments. Significant differences were determined using an unpaired two-tailed *t* test (*p*= 0.018). H and I) Image and plot represents the invasion properties of WT and N106S mutant as determined using a colorimetric transwell invasion assay using 10% serum as the chemoattractant. Colorimetric measurements of cell invasion were plotted as a percent of control cells plated with no chemoattractant. Plot indicates standard error of the mean from three independent experiments. Significant differences were determined using an unpaired two-tailed *t* test (*p*= 0.031).

The TEs that were downregulated showed no correlation with the length of 5’UTR and only showed mild correlation with the length of coding region (Fig. 7C and S6B). On the other hand, TEs of transcripts with long 5’UTRs, higher GC content and long coding regions were upregulated in the mutant (Fig. 7E, 7F and S6D). Interestingly, transcripts with high GC content and long 5’UTRs are characteristic of mRNAs that are often challenging to translate and prefer cap-independent translation suggesting a potential shift in cap-dependent to cap-independent translation as seen during mitosis. However, there was no correlation with 3’UTR length for upregulated TEs (Fig. 7D) or with the overall transcript length (Fig. S6C and E).

### eIF6-N106S mutation markedly affects colonic cancer cell invasion properties in vitro and in vivo

Ribo-seq analysis also showed significant alterations in TEs of cytoskeletal factors especially downregulation of microtubule-associated genes and upregulation of several other cytoskeletal factors. (Fig. 6D and E). Given that Tubulin staining showed altered cell to cell contacts, and given the importance of eIF6 in cancer progression, we wanted to determine the effects of eIF6-N106S mutation on the invasive properties of HCT116 cells. HCT116 cells are a highly invasive colonic carcinoma cell line that are known to metastasize in mice xenograft models^49^. We therefore tested the cell migration and invasion properties of the eIF6-N106S mutant cells both *in vitro* and *in vivo*. eIF6-N106S mutant cells showed limited cell migration abilities compared to WT in a transwell migration assay *in vitro* (Fig. 7G). eIF6-N106S mutant cells also showed a lack of ability to invade through the extracellular matrix in the transwell cell invasion assay (Fig. 7H and I).

HCT116 colonic cancer cells have unique invasive properties in that they are highly invasive but show poor epithelial to mesenchymal transitions (EMT) and have retained their epithelial phenotype ^50–52^. They show characteristics of cancer cells capable of invasion through collective cell migration with minimal EMT changes^53^. However, to better understand the changes in invasion properties of mutants, we probed for changes in markers of epithelial to mesenchymal transition (EMT): Snail, Slug, and E-Cadherin (Fig. S8A and B). We did not observe any changes in the EMT markers between WT and mutant. Previous studies have shown that overexpression of eIF6 in ovarian cancers and melanomas that already have high levels of eIF6 promote invasion by increasing the protein levels of Cdc42^54^. However, we did not observe any changes in total Cdc42 levels in mutants compared to WT (Fig. S8B). To better understand the effects of eIF6-mutation in limiting cancer invasion, we analyzed the Ribo-Seq data and found that the translation efficiencies of several factors that are critical for metastasis of many cancers such as nuclear factor I/B (NFIB)^55,56^, and vasodilator-stimulated phosphoprotein (VASP)^57–59^ were markedly downregulated in the mutant cells (Fig. 7J). We therefore tested for changes in expression of these genes. NFIB has many isoforms, but the predominant and most-studied isoform is about 47kDa. We found that the expression of the predominant isoform was almost absent in the eIF6-N106S mutant (Fig. 7J). VASP expression was also absent in the eIF6-N106S mutant cells (Fig. 7J). This indicates a downregulation of specific factors critical for metastasis.

Given the effects of eIF6-N106S mutation on cancer invasion properties *in vitro*, we wanted to determine the effect of mutation on cancer growth and invasion *in vivo* using a tumor xenograft model. Both eIF6-WT and eIF6-N106S mutant cells were stably transfected with firefly luciferase to monitor tumor growth and invasion *in vivo* (Fig. 8A). To avoid signal bias, clones with similar bioluminescence signal and firefly luciferase activity were selected (Fig. 8A and B). Clones were also screened by morphological assessment and cell proliferation assay to ensure that they have retained the properties of parental clones (Fig. 8C). We then subcutaneously injected 7-to 8-week-old NOD-Scid immunocompromised mice with eIF6-WT cells (n=8 mice) or eIF6-N106S cells (n=8 mice). The luciferase activity on day 0 was found to be similar between WT and the mutant mice indicating that any difference between WT and mutants was not due to variable cell numbers during injections (Fig. 8D).

**Figure 8.**
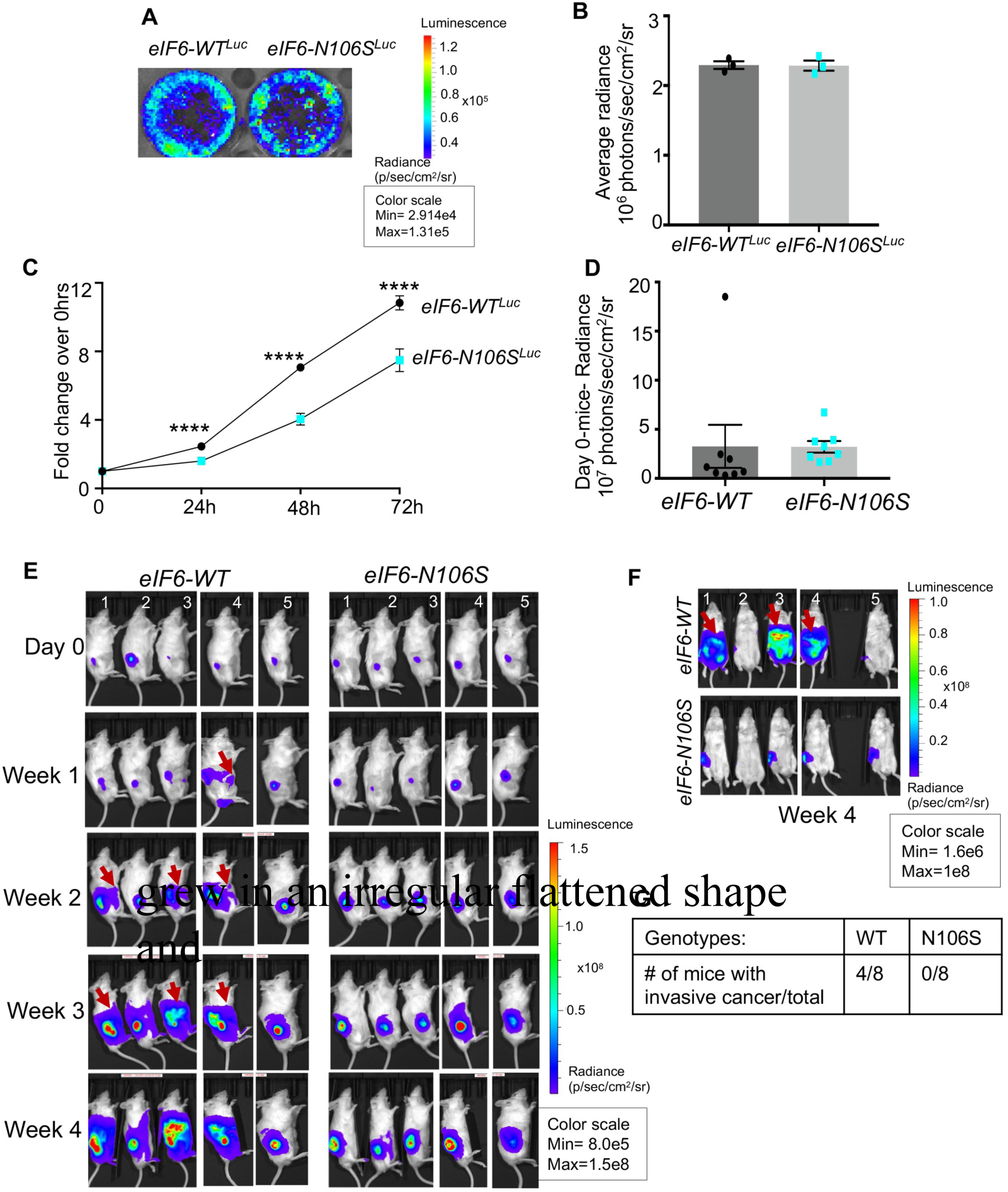
eIF6-N106S mutation inhibits colonic cancer invasion in a subcutaneous tumor xenograft model. A. Bioluminescence image shows stable transfectant clones of eIF6-WT and eIF6-N106S cells expressing similar firefly luciferase activity. B. Luciferase activity for images shown in A. were quantitated and plotted as average radiance. Plot indicates standard error of the mean of three independent experiments. The difference between eIF6-WT and eIF6-N106S was not significant (*p*=0.94) as determined by an unpaired two-tailed *t* test. C) Plot indicates the fold change in cell proliferation rates relative to 0 hours of plating as measured by the MTS assay. Plot indicates standard error of the mean of three independent experiments with triplicate wells measured per experiment. The difference between eIF6-WT and eIF6-N106S was significant at 24 hours (*p*=0.000001), 48 hours (*p*=0.000001) and 72 hours (*p*=0.0005) of growth as determined by an unpaired two-tailed *t* test. D) Plot indicates the average luciferase activity in mice injected with eIF6-WT cells (n=8 mice) and eIF6-N106S cells (n=8) on the day of injection (Day 0). The difference between eIF6-WT and eIF6-N106S was not significant (*p*=0.98) as determined by an unpaired two-tailed *t* test. E) Representative images show bioluminescence signal measured from Day 0 to week 4 before the mice were sacrificed. Photon flux is indicated by pseudocolored heatmap and tumor-specific luciferase activity was measured by defining region of interest (ROI). The presence of luciferase activity in other parts of the body beyond the subcutaneous injection site indicated with red arrows. F) Representative images of ventral view of mice showing bioluminescence signal at week 4. G) Table indicates the number of mice exhibiting invasive cancer relative to total number of mice.

The eIF6-WT cells as expected were highly invasive and four out of eight mice showed the tumor invading through the subcutaneous layer and protruding into the peritoneal space and three of the mice showed extensive peritoneal metastases and hepatic metastases (Fig. 8E and S9A and B). The metastatic rates in the subcutaneous model can be variable and so, we observed invasion in 50% of the eIF6-WT tumors but none of the eIF6-N106S tumors showed invasion. The eIF6-N106S primary tumor growth was slower compared to eIF6-WT tumors and as shown by increased fold-change in luciferase activity in WT tumors (Fig. S9C). However, the dome-shaped subcutaneous primary tumor sizes could not be accurately determined in WT as half of the WT tumors grew inwards due to invasion through the subcutaneous layer (Fig. S9A and S9D). The external tumor shape was also variable and often displayed flattened tumors (Fig. S9A). However, even by day 28, when the tumor size of eIF6-N106S was large and mice had to be sacrificed due to weight loss, none of the eIF6-N106S tumors showed any invasion. These results indicate that the eIF6-N106S mutation markedly inhibits the cancer invasive properties of colonic cancer cells even *in vivo*.

### eIF6 is overexpressed primarily in high-grade invasive cancers

Given the importance of eIF6 for invasion, we wanted to examine the correlation between eIF6 expression and cancer invasiveness using bladder cancer patient samples and breast cancer cell lines. These two tissue types were specifically selected as eIF6 role in bladder and breast cancers is poorly understood. In addition, majority of cancer studies on eIF6 have thus far shown high eIF6 expression in cancers in comparison to non-neoplastic tissues^6^. However, very few studies have examined if overexpression of eIF6 is universal to the cancerous state regardless of subtypes and grades^31,60^. To better understand changes in eIF6 expression, we screened a panel of Grade 3 and Grade 4 bladder cancer cell lines (Fig. 9A and B). Results show that eIF6 is expressed in many high-grade bladder cancer cell lines relative to the healthy control primary bladder cells (Fig. 9A and B). However, eIF6 was not highly expressed in all cancer subtypes as suggested previously.

**Figure 9.**
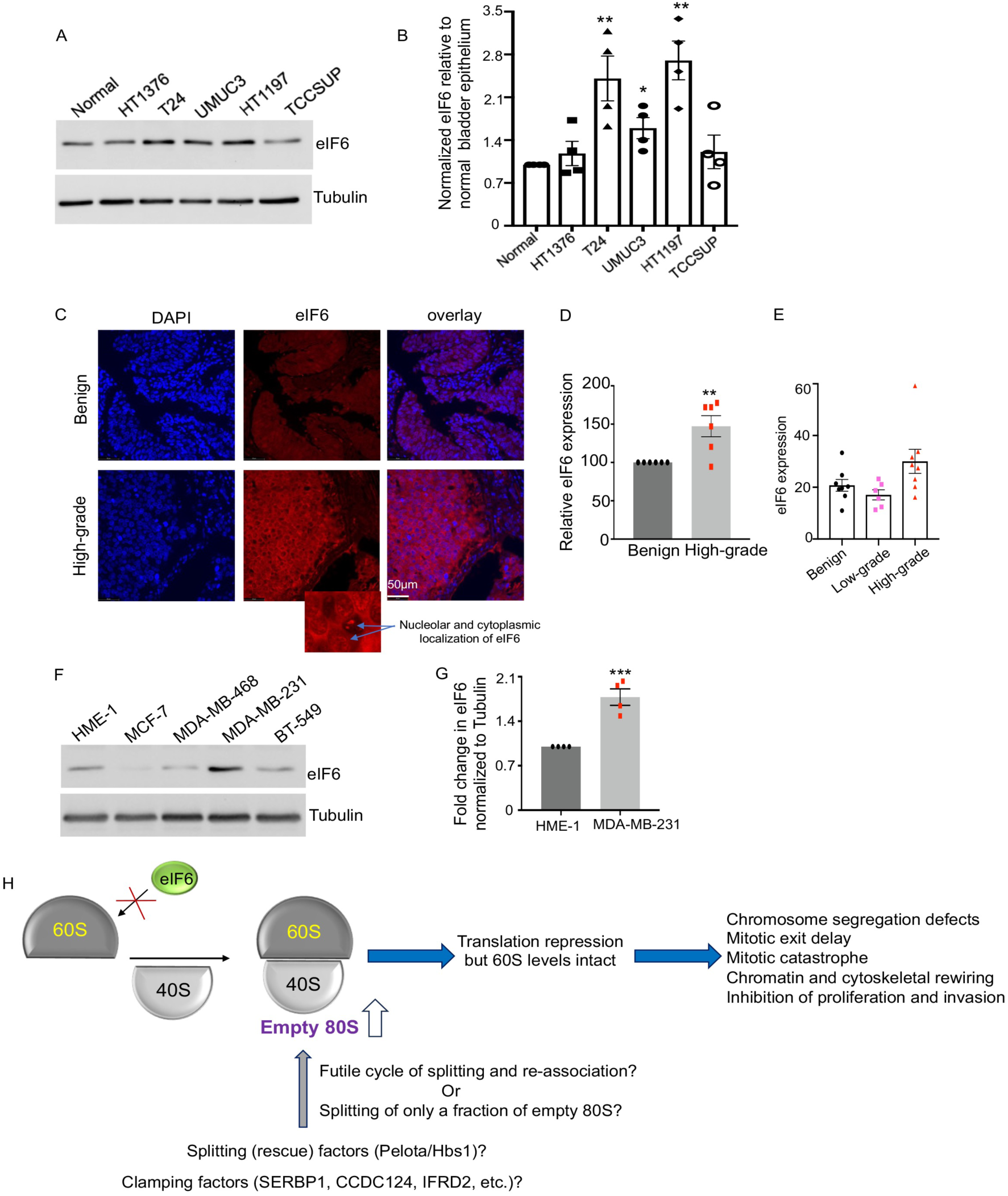
Overexpression of eIF6 observed primarily in high-grade invasive bladder and breast cancers. A and B) Western blot probed for eIF6 levels in human bladder cancer cell lines and healthy (normal) bladder epithelial cells. β-Tubulin used as loading control. Blots shown in A. were quantitated and eIF6 levels were normalized to loading control and plotted (B) as fold change over eIF6 levels in normal bladder epithelial cells. Values indicate standard error of the mean from four independent experiments and significant differences for T24 (*p*=0.0065), UMUC3 (*p*= 0.0132) and HT1197 (*p*=0.0017) determined by an unpaired two-tailed *t* test. C) Images represent eIF6 protein expression in patient-derived benign and high-grade tumors by immunohistochemistry using anti-eIF6 antibody. Enlarged inset shows the presence of eIF6 in nucleoli and cytoplasmic localization of eIF6 in high-grade cancers. D) Images shown in C. were quantitated and eIF6 expression in patient-matched benign tissues relative to high-grade cancers (6 patients) were plotted. Significant differences were determined using an unpaired two-tailed *t* test. (*p*=0.0063). E) Plot shows eIF6 expression in un-matched benign tissues, high-grade and low-grade cancers derived from patients. F and G) Western blot represents high levels of eIF6 in high-grade invasive human triple negative breast cancer cell line. β-Tubulin was used as loading control. Blots shown in E. were quantitated and eIF6 levels were normalized to loading control and plotted (B) as fold change over eIF6 levels in normal (healthy) HME-1 cells. Values indicate standard error of the mean from four independent experiments and significant differences for MDA-MB-231 (*p*=0.001) determined by an unpaired two-tailed *t* test. H. Model depicts the effects of disrupting eIF6 interaction with the 60S subunit using the eIF6-N106S mutation. Model is detailed in discussion.

We also screened for changes in eIF6 expression using a panel of bladder cancer patient samples where the benign tissue of the patient was matched to the high-grade cancers (Fig. 9C and D). Tissues stained with secondary antibody-only control was used to confirm the specificity of staining (Fig. S10). eIF6 was overexpressed only in the high-grade cancers and eIF6 expression was predominantly localized to the cytoplasm and nucleoli (Fig. 9C and D). However, we did not observe any changes in eIF6 expression in low-grade bladder cancers relative to the benign tissues (Fig. 9E). This suggested that eIF6 is not overexpressed in all bladder cancers alike. A previous genomics study had uncovered gene amplification of eIF6 in breast cancers and they predicted eIF6 to be an early driver of breast cancers^61^. However, eIF6 expression in breast cancers is yet to be examined. We therefore screened a panel of breast cancer cell lines of different subtypes relative to the healthy breast epithelial cell line (HME-1) (Fig. 9F). eIF6 expression was found to be highly induced only in the highly invasive and high-grade triple negative breast cancer cell line: MDA-MB-231 (Fig. 9F and G). eIF6 expression was not induced in other breast cancer cell lines. These results suggests that eIF6 is not overexpressed in all cancers alike, instead, it is preferentially induced only in highly invasive and high-grade cancers. It also indicates that induction of eIF6 is not an early event in breast and bladder cancers.

## Discussion

In this study, we modulated the pools of active 80S that are available to engage in translation by using the eIF6-N106S mutation that severely weakened eIF6 interaction with the 60S subunit. This effect of N106S mutation in disrupting 60S binding was also shown in *Dictyostelium* cells where the N106S mutant was expressed in cells lacking endogenous eIF6. It is intriguing that even though we severely disrupted the anti-association activity of eIF6 that led to an increase in vacant 80S levels, there was still a pool of 80S available to engage in translation and to maintain a basal level of protein synthesis that allowed cells to continue to divide albeit under a stressed state. This suggests that there may be compensatory mechanisms to rescue some of the vacant 80S ribosomes and permit their re-engagement in translation. However, these mechanisms are unable to fully rescue the vacant 80S pools.

The vacant (empty or idle) 80S mammalian ribosomes exist in distinct classes defined by the type of clamping factors that stabilize the vacant 80S^62^. The predominant class of vacant 80S ribosomes are stabilized by Serpine mRNA binding protein (SERBP1) (Stm1 in yeast) and eEF2^62–65^. Another minor class of vacant 80S ribosomes are stabilized by binding to coiled-coil domain containing 124 protein (CCDC124) (Lso2 in yeast)^62,64,66^. Mammalian rescue factors: Pelota/Hbs1/ABCE1 and yeast rescue factors: Dom34/Hbs1/Rli1 can efficiently split the vacant 80S ribosomes^62,67,68^. *In vitro*, the Stm1-bound ribosomes are more resistant to splitting by the Dom34 (Pelota)-Hbs1 rescue factors as opposed to the Lso2-bound ribosomes^64^. Therefore, it is possible that in the eIF6-N106S mutants, only a small fraction of vacant 80S ribosomes that are much conducive to splitting are split by the rescue factors but a major fraction of vacant 80S is not efficiently dissociated. Alternatively, it is likely that the vacant 80S ribosomes are split in the mutants, but in the absence of eIF6 to inhibit association, they go through a futile cycle of splitting and re-association that eventually limits the pool of active 80S (Fig. S9H, model). Future studies on the composition of the 80S in the mutants will help to address the mechanism involved.

Another intriguing finding in this study is the effect of limiting ribosomal availability in disrupting the mitotic phase of cell cycle much more than other phases of cell cycle. This specific effect of hindering mitotic progression by limiting ribosomal availability has been previously observed for the rescue factor-Pelota^69^. Loss of Pelota in mouse embryos affects mitotic progression leading to aneuploidy and cell death^69^. We observe a similar effect in the eIF6-N106S mutant, specifically in delaying mitotic exit and cytokinesis. This suggests that active 80S ribosomal availability is rate limiting for the recovery from translational inhibition that is needed to exit mitosis. It also suggests that limiting ribosomal availability up to a certain threshold can be tolerated by cells and they can survive and cycle through as long as the global translation rates are largely undisturbed. However, when cells experience conditions in which global translation rates fluctuate with an inhibition phase followed by recovery (as seen in mitosis or as seen during stressed states such as nutrient deprivation)^36–43^, then limiting ribosomal availability can become detrimental to cells. This has been shown by the critical functions of rescue factors (Dom34 (Pelota)/Hbs1) and clamping factors such as Stm1p and Lso2p in aiding cells recover from nutrient stress conditions^65,66,68^. We also observed this previously using a C-tail mutant of eIF6, where the translation inhibition observed in the eIF6-mutant cells was shown to be important for cells to adapt to starvation^70^. Also, eIF6 ^+/-^ heterozygous knockout mouse embryonic fibroblasts showed delayed S-phase upon recovery from serum starvation^7^. Whether altering mitotic progression and stress response by modulating the stoichiometry of active ribosomal subunits contributes to the genomic instability and cancer-predisposition of certain ribosomopathies remains to be further explored.

The 60S-interaction site mutant (N106S) of eIF6 reveals a dichotomy in the anti-association function of eIF6 and its role in nucleolar 60S biogenesis. Previous studies in yeast showed that depleting eIF6, especially in the nucleolus, leads to cell death by causing a marked loss of total 60S levels due to defects in pre-rRNA processing^10^. With the N106S mutation, even though we markedly disrupted eIF6 interaction with 60S, we did not observe marked changes in the total levels of 60S or in the nucleolar localization of eIF6. This suggests that eIF6 function in 60S biogenesis is well preserved even when eIF6-60S interactions are disrupted. This has been previously observed in the *eIF6* heterozygous knockout mice where partial loss of eIF6 markedly affected its anti-association activity without disrupting 60S levels^7,18^. This suggests that targeting the uL14 (RPL23) interface of interaction of eIF6 will not adversely alter 60S levels and so, this interface could be therapeutically targeted in cancers and Shwachman-Diamond syndrome with potentially minimal side effects. It also suggests that the N106 interaction-interface is a desirable target. It is possible that the effect on cancer growth by disrupting the N106 residue is much more severe than disrupting other residues in the interaction interface. This is supported by previous studies showing that among the compensatory somatic mutations in eIF6 seen in SDS patients, the N106S mutation is favored much more than other interaction-site mutations in eIF6.

The nucleolar localization of eIF6 was previously shown to be dependent on direct interactions of eIF6 with BCCIP^35^. However, two other studies using yeast Tif6 and human eIF6 showed that the interaction of eIF6 with BCCIP are indirect and mediated only through eIF6 interactions with the uL14-BCCIP complex^71,72^. Our study supports these latter two studies and indicate an indirect interaction of eIF6 with BCCIP, since abrogating eIF6 interaction with uL14 also disrupted its interaction with BCCIP, but it did not disrupt the nucleolar localization of eIF6. This suggests that eIF6 interaction with BCCIP is not critical for its nucleolar localization. It is possible that another redundant pathway might compensate for the loss of uL14-BCCIP-mediated nucleolar recruitment of eIF6.

In this study, we also observed changes in two key regulators of invasion in mutants: VASP and NFIB. NFIB has been shown to promote invasion through increasing chromatin accessibility^55,56^. Our Ribo-Seq data shows major upregulation of histone proteins in the mutants suggesting that there could be an increase in nucleosomes that could make the chromatin less accessible. It remains to be determined if the chromatin state is altered in the mutants and if rescuing NFIB can restore the chromatin state and enhance invasion in the mutant cells. Although, given the extent of translational deregulation, rescuing just one or two factors may not fully revert the phenotypic effect of mutation. Also, loss of VASP inhibits invasion by altering the lamellipodia architecture^57–59^. Preliminary morphological analysis of the mutant cells showed a decrease in lamellipodia. Future studies will determine the effects of eIF6 mutation on lamellipodia organization and invasion.

Our analysis of bladder and breast cancer cells along with few other previous studies especially in colon cancers suggest that eIF6 expression is highly induced in the invasive stages of cancer progression^31,60^. This suggests that induction of eIF6 is less likely to be an early event that drives tumorigenesis. This is supported by a recent study showing that the inducible overexpression of eIF6 in normal c-Kit+ bone marrow cells is not conducive for growth as it inhibits global translation by causing a subunit joining defect^5^. This suggests that overexpression of eIF6 is not sufficient to drive tumor growth by itself. However, as the tumor progresses, the enhanced translational demands and high ribosomal levels could induce a compensatory increase in eIF6 expression to ensure that an appropriate stoichiometry of active subunits are made available to engage in translation in high-grade invasive cancers. Future studies using a metastatic model of tumor progression are needed to better understand the role and mechanism of eIF6 induction in invasive cancers.

## Supporting information

All Suppl. images

## Acknowledgements

We thank the RNA Core at Case Western University, Dr. Andrew Hsieh at Fred Hutchinson Cancer Center for his helpful comments, Dr. Miao Yong at the Genome Engineering and IPSC Center at Washington University in St. Louis, Mr. Joseph Loredo and Dr. Daniela Masson-Meyers for their technical support. We also thank Ms. Melanie Wiese, Ms. Jenifer Sterling, Ms. Cynthia Kurklis, Dr. John Long and Dr. Kathleen Donovan in the Division of Comparative Medicine for their technical assistance with the mice xenograft studies.

## Data Availability

All data have been included in the manuscript.

## Funding and Additional Information

This work was supported by grants from the National Institutes of Health R01GM143179 to Sofia Origanti and National Science Foundation #1951332 to Zhenguo Lin.

## Conflict of Interest

None

## Materials and Methods

### Cell culture and reagents

All cell lines used in this study were obtained from ATCC. The human colorectal carcinoma line (HCT116) was maintained in McCoy’s 5A (modified) medium with 10% fetal bovine serum (FBS) and 100 units/mL penicillin and 100 mg/mL streptomycin (Gibco) (Penn/Strep). The generation of *eIF6^N^*^106^*^S/N^*^106^*^S^* homozygous mutant clones by CRISPR/Cas9 editing and their screening by next-generation sequencing and STR analysis was described in our previous study^30^. Clones are not passaged for more than six to seven passages. The human breast cancer cell lines MDA-MB-468, MDA-MB-231 were maintained in DMEM medium with 10% FBS and Penn/Strep, BT-549 was maintained in RPMI-1640 with 10% FBS and Penn/Strep, MCF-7 was maintained in EMEM medium with 10% FBS and Penn/Strep, normal human breast cell line (hTERT-HME-1) was maintained in DMEM F12 media with 10% FBS and Penn/Strep. The human bladder cancer cell lines HT-1376, HT1197, UM-UC-3 and TCCSUP were maintained in EMEM media with 10% FBS and Penn/Strep, T-24 cell lines were maintained in McCoy’s 5A medium with 10% FBS and Penn/Strep, primary bladder epithelial (A/T/N): normal human (BdEC) cells were maintained in bladder epithelial cell basal medium (ATCC) supplemented with bladder epithelial growth factors. MTS assay for the second clone of eIF6-N106S homozygous knock-in HCT116 cells was performed as described before^30^. Cells are routinely tested using mycoplasma PCR detection kit (ATCC).

### Western blot analysis

Cells were lysed in Mammalian Cell Lysis Buffer (MCLB) (50 mM Tris-Cl pH 8.0, 5 mM EDTA, 0.5% IGEPAL, 150 mM NaCl) supplemented with the following inhibitors right before lysis: 1 mM phenylmethylsufonyl fluoride (PMSF), 1 mM sodium fluoride, 10 mM β-glycerophosphate, 1 mM sodium vanadate, 2 mM DTT, 1X protease inhibitor cocktail (Sigma–Aldrich), 1X phosphatase inhibitor cocktail (Santa Cruz Biotechnology). Cell lysates were rocked for 15min at 4°C and centrifuged at 14,000 RPM for 10min at 4°C. Lysates were resolved by SDS-PAGE and transferred onto nitrocellulose membrane (0.45µm; Bio-Rad Laboratories). Blots were blocked for 1 hour at room temperature in 5% nonfat dry milk dissolved in Tris Buffered Saline with 0.1% Tween 20 (TBS-T). The following antibodies were diluted in TBS-T buffer: eIF6 (1:1000; Santa Cruz Biotechnology), eIF6 (1:1000; Cell Signaling), RPL23 (1:2500, Bethyl), β-Tubulin (1:2500; Cell Signaling), GAPDH (1:6000, Cell Signaling), Vinculin (1:1000; Cell Signaling), β-Actin (1:10000, Cell Signaling), p53 (1:1000; Santa Cruz Biotechnology), p21 (1:1000; Santa Cruz Biotechnology), BCCIP (1:1500; Bethyl), eIF4E (1:1000, Cell Signaling), 4EBP1 (1:1000, Cell Signaling), 4EBP2 (1:1000, Cell Signaling), eIF2α (1:1000, Cell Signaling), eIF2A (1:1000, Bethyl), eIF4G1 (1:1000, Cell Signaling), topoisomerase IIβ (1:1000, BD Biosciences), eIF4G2 (1:4000, Bethyl), SBDS (1: 1000, Cell Signaling), VASP (1:1000, Cell Signaling), Cdc42 (1:1000, Cell Signaling), β-Catenin (1:1000; Santa Cruz Biotechnology), E-Cadherin (1:1000; Santa Cruz Biotechnology), NFIB (1:2000 in 2% milk, Bethyl), Snail (1:1000; Cell Signaling). All phospho-specific antibodies were diluted in TBS-T with 1% milk and following antibodies were used; phospho-Ser-209-eIF4E (1:1000; Cell Signaling), phospho-Ser-51-eIF2α (1:1000, Cell Signaling), SLUG (1:1000, Cell Signaling), EFL1 (1:2500, Bethyl), phospho-Ser-10-Histone H3 (1:1000, Cell Signaling), and anti-γH2AX (1:1000; Cell Signaling). Membranes were probed with HRP-conjugated secondary antibodies 1:30,000 diluted in TBS-T buffer (Jackson Immunoresearch). To induce eIF2α phosphorylation by oxidative stress, cells were treated with 4 mM H_2_O_2_ for 1 hour. Blots were developed with ECL substrate (Pierce) and images were acquired in the iBright CL1500 imager (Thermo Fisher Scientific). Blots were quantitated using iBright analysis software. For analysis of phospho-Ser-10-Histone H3 levels, cells were synchronized in mitosis with 75 ng/mL nocodazole (Sigma-Aldrich) for 18 hours. Mitotic cells were collected by shaking off rounded cells as described before^73^. The eIF6-N106S mutant is thermally unstable *in vitro* as shown in our previous CD analysis^30^ and therefore, the mutant is prone to mild degradation *in vitro* with increasing number of freeze-thaws of cell lysates and based on the lysis buffer used.

### Click-iT translation assays

Global translation rates were measured using Click-iT AHA (L-Azidohomoalanine) Alexa Fluor 488 Protein Synthesis HCS assay kit as per manufacturer’s instructions (Invitrogen). Briefly, 7,000 HCT116 cells were seeded per well of 96-well plates and cultured for 48 hours. After 48 hours, cells were washed twice with 1X phosphate buffered saline (PBS) and incubated in methionine-free media and Click-iT AHA reagent (50 µM) or DMSO control for 30 min at 37°C in a CO_2_ incubator. The cells were then washed and fixed in 4% paraformaldehyde (PFA) for 40 min. Azido-modified proteins were detected by cycloaddition reaction with an Alexa Fluor 488-conjugated alkyne, and nuclei were stained with Hoechst 33342 (10 µg/mL) to determine cell number per well. Fluorescence intensities were recorded using a microplate reader (Synergy H1, BioTek) and fluorescence intensity in each well was normalized to nuclear fluorescence intensity. The translation rates were calculated by normalizing to DMSO control.

Translation rate in mitosis were estimated using Click-iT OPP (O-propargyl-puromycin) Alexa Fluor 488 Protein Synthesis HCS assay kit (Invitrogen). Briefly, 5,000 cells were cultured in 96-well plates for 48 hours, followed by addition of Nocodazole (75ng/mL) (Sigma) for 18 hours to synchronize cells in mitosis. For the asynchronous HCT116 cells, 3,000 to 4,000 cells were seeded in 96-well plates for 72 hours. The cells were then incubated with Click-iT OPP reagent (20 µM) for 30 min at 37°C. The cells were fixed in 4% PFA for 1 hour, and alkyne-modified proteins were detected with an Alexa Fluor 488 picolyl azide, and nuclei were stained with HCS nuclear mask Blue stain to determine cell number in each well. Fluorescence intensities were measured and translation rates were calculated as described above.

### Immunofluorescence

HCT116 cells were cultured on poly-D-lysine–coated glass coverslips (Neuvitro) for 48 hours and washed once with 1X PBS prior to fixation. Cells were fixed in 4% PFA for overnight at 4°C. Cells were permeabilized for 15 min using 2% TritonX-100 diluted in PBS and incubated in blocking buffer (2% Bovine Serum Albumin -BSA in 1X PBS and 0.1% IGEPAL) for 30 min at RT. For immunostaining, α-Tubulin (1:45; Cell Signalling) was diluted in blocking buffer and incubated for 1 hour at RT. For analysis of eIF6 localization, coverslips were incubated with anti-eIF6 antibody (1:100; Santa Cruz Biotechnology) diluted in blocking buffer and incubated for 1 hour at RT. Coverslips were washed three times in wash buffer (1X PBS, 0.1% IGEPAL) and incubated for 45 mins with either donkey anti-rabbit Alexa Fluor 546 (1:500), donkey anti-mouse Alexa Fluor 546 (1:500) or goat anti-rabbit Alexa Fluor 488 (1:500) secondary antibodies (Thermo Fisher Scientific). For staining with anti-centromere antibody, asynchronous cells were fixed with 3.5% paraformaldehyde for 15 min at RT, washed twice with 1X PBS and then fixed in ice cold 100% methanol for 10 min at 4°C. Cells were permeabilized with 0.2% Triton X-100 in 1XPBS for 15 min at RT. Cells were then incubated with blocking buffer (3% BSA in 1X PBS) for 30 min at RT. After washing with 1X PBS three times, cells were incubated with α-Tubulin (1:45; Cell Signaling) diluted in blocking buffer and incubated for 1 hour at RT. After washing with 1X PBS, coverslips were stained with human anti-centromere (CREST) antibody (1:1000, Immunovision) diluted in blocking buffer. Coverslips were then washed again and incubated with secondary antibody Goat anti-human Alexa Fluor 647 (1:500) and diluted in blocking buffer B for 45 min at RT. Coverslips were rinsed four times with wash buffer and mounted using ProLong Gold antifade reagent with DAPI (Invitrogen). Immunofluorescence was analyzed under Leica DM6 B upright fluorescent microscope and digital images were captured through Leica K8 CMOS camera (Leica) using LAS X imaging software. LUT is linear and covers the full data range. A total of about 900 asynchronous cells from three independent experiments were analysed to calculate the percentage of cells exhibiting defects in chromosome segregation, cytokinesis defects and multinucleation.

### Time-lapse live imaging

Cells were cultured in 35 mm glass bottom dishes (MatTek) and treated with 10 µM RO-3306 (CDK1 inhibitor) (Tocris) for 20 hours. Drug was removed by washing the cells twice with 1X PBS and fresh culture medium was added. Live cell-time lapse imaging was performed using SP8 confocal microscope (Leica) with 20X objective (Leica) and images were captured using DFC7000GT camera. Cells were placed on a microscope stage incubator (Okolab) set at 37°C with 5% CO_2._ Images were captured at 2 min 30 sec intervals for 3 hours and processed using LAS X software (Leica).

### Immunoprecipitation

Cells were lysed in cold MCLB buffer supplemented with inhibitors as described above. For immunoprecipitation (IP) of endogenous eIF6, 1000 µg of total protein suspended in 500 µL of cell lysate was precleared with 20 µL of normal mouse IgG-conjugated agarose beads (Santa Cruz Biotechnology; sc-2343) for 30 mins at 4°C on a rotator. The precleared lysates were incubated with 10 µL of anti-eIF6 antibody conjugated agarose beads (Santa Cruz Biotechnology; 390432) for overnight at 4°C on a rotator. For IP of endogenous BCCIP, 1000 µg of total protein suspended in 500µL of cell lysate was precleared with 20 µL of protein A/G plus agarose beads (Santa Cruz Biotechnology) and 1 µL of normal rabbit IgG (Cell Signalling; 2729). The precleared lysates were incubated with 20 µL of protein A/G plus agarose beads and 2 µg of BCCIP antibody (Bethyl; A302-196A). Beads were washed three times in cold MCLB buffer and resuspended in 30 µL of MCLB buffer. Samples were eluted by boiling for 15 min in 1X Laemmli SDS buffer with DTT and analyzed by western blotting.

### Cell migration and invasion assay

Transwell migration assay was performed using Chemicon QCM^TM^ 24-well kit (Millipore, ECM509). 1.5 x 10^6^ cells were seeded on 100-mm plates and serum starved for 18 to 20 hours and harvested in McCoy’s media with 5% BSA, and 250,000 cell suspension in 300 µL volume was plated onto 8 µm pore size inserts and placed on top of the lower chamber with media that had either 10% FBS (chemoattractant) or no-serum control. Plates were incubated for 24 hours at 37°C and the cells that migrated were collected in cell detachment solution, followed by incubation with CyQUANT GR dye. Fluorescence intensities were recorded on microplate reader (Synergy H1, BioTek) and relative fluorescence units (RFU) were calculated by subtracting control values for the respective genotype.

Transwell invasion assay was performed using colorimetric cell invasion kit (Chemicon, Millipore; ECM550). Cells were serum starved and extracellular matrix (ECM) coated inserts were rehydrated for 1 hour with media without serum. 250,000 cell suspension in 300 µL volume was plated onto 8 µm ECM-coated insert and placed on top of the lower chamber with media that had either 10% FBS (chemoattractant) or without serum (control) and plates were incubated for 48 hours at 37°C. Cells that invaded were stained with crystal violet stain (Millipore, #20294) and intensity of staining was analysed by ImageJ software to determine the area of invasion.

### Ethics statement

All animal experiments in this study were conducted according to the approval (protocol #2992) and guidelines of Saint Louis University Institutional animal care and committee (IACUC).

### Subcutaneous injections and bioluminescence imaging

HCT116 cells (eIF6-WT and eIF6-N106S) were transfected with 4µg of pcDNA-firefly-luciferase plasmid DNA (Addgene; 18964) and 20 µL of Lipofectamine 2000 (Invitrogen). 24 hours post transfection, cells were selected in medium containing 500 µg/mL G418 (Gibco). About 10 drug resistant colonies from each cell line were selected and maintained in culture media containing 100 µg/mL G418. Stable transfectants were screened for luciferase activity by adding 300 µg/mL D-luciferin (Gold Biotechnology) and bioluminescence was captured using IVIS Lumina (Revvity). Stable cell lines with similar luciferase expression were selected. 7 to 8 weeks old (NSG) NOD.Cg-Prkdcscid Il2rgtm1Wjl/SzJ (Jackson Laboratory) male mice weighing 24 g to 28 g were subcutaneously injected in the right flank with 1.5 x 10^6^ cells suspended in 100 µL of McCoy’s media. Injections were performed in two batches of 5 (WT/N106S) mice and 3 mice (WT/N106S) respectively by two different veterinary assistants at Saint Louis University, who were blind to the study. For imaging tumors *in vivo*, mice were injected with 150 mg/kg D-luciferin (Gold Biotechnology) intraperitoneally and bioluminescence was acquired using IVIS Lumina (Revvity) using the following acquisition parameters: exposure time-5 min, Binning-8, no filter; f/stop, FOV-12.5cm. Photon flux is indicated by pseudocolored heatmap and tumor-specific luciferase activity was measured (units: photons/sec x 10^5^ to 10^8^) by drawing a circular region of interest (ROI). For measurement of primary tumors, once the tumor became visible at 14 days, the length and width of tumor was measured each week with a digital caliper (Mitutoyo) and tumor volumes were calculated using the formula: (LxWxW)/2. Mice were euthanized as per IACUC guidelines and humane endpoints were followed as per IACUC guidelines.

### MTS assay

Viability of eIF6-WT^Luc^ and eIF6-N106S^Luc^ cells was performed using MTS assay. 5,000 HCT116 cells expressing eIF6-WT or eIF6-N106S mutant was cultured per well of a 96-well plate in McCoy’s culture medium without phenol red supplemented with 10% FBS and Penn/Strep. 20 μL of MTS solution (CellTiter 96^®^ Aqueous One solution reagent, Promega) was added to each well, and the plates were incubated for 1 hour. Absorbances were read at 490 nm (Synergy H1, BioTek). To obtain background-corrected absorbances, average absorbance values of the control wells containing media and MTS only were subtracted from all other absorbances.

### Subcellular fractionation

700,000 HCT116 cells were cultured in 100-mm dish for 48 hours. To assay for total protein input, cells were lysed in MCLB buffer supplemented with inhibitors as previously described^70^. To extract cytoplasmic fractions, the cells were collected in cell lysis buffer (10 mM HEPES, pH 7.5, 10 mM KCl, 0.1 mM EDTA, 0.5% IGEPAL) supplemented with the following inhibitors: 10 mM β-glycerophosphate, 1 mM sodium vanadate, 2 mM DTT, 1X protease inhibitor cocktail (Sigma–Aldrich), 1X phosphatase inhibitor cocktail (Santa Cruz Biotechnology), and 0.5 mM PMSF. For cytoplasmic fractionation, the lysates were incubated on ice for 15 min with intermittent gentle mixing by inverting the tube and vortexing on high for 10 sec. The cells were centrifuged at 12,000xg for 10 min at 4 °C, and the cytoplasmic supernatant fraction was collected. For nuclear extraction, nuclear pellets were washed four times in cell lysis buffer and centrifuged at 3,000 RPM for 5 min at 4 °C. Nuclear pellets were suspended in 75 μL of nuclear extract buffer (20 mM HEPES, pH 7.5, 400 mM sodium chloride, 1 mM EDTA) supplemented with the following inhibitors: 10 mM β-glycerophosphate, 1 mM sodium vanadate, 2 mM DTT, 1X protease inhibitor cocktail (Sigma–Aldrich), 1X phosphatase inhibitor cocktail (Santa Cruz Biotechnology), and 0.5 mM PMSF. Nuclear pellets were solubilized on ice for 30 min by vortexing on high every 10 min for 15 sec followed by centrifugation at 12,000xg for 15 min at 4 °C. Both nuclear and cytoplasmic fractions were analyzed by Western blotting.

### Polysome profile analysis

Polysome profile analysis was carried out as described previously^70^, except 1.5 million cells for eIF6-WT, and 2 million cells for eIF6-N106S (50% to 60% cell density) were plated per 100-mm dish. The cells were treated with 100 µg/mL cycloheximide (Tocris) for 5 min at 37 °C and washed with ice-cold 1X PBS buffer containing 100 µg/ml cycloheximide on ice. Cells were lysed in 1X polysome lysis buffer (50 mM HEPES pH 7.4, 250 mM KCl, 5 mM MgCl_2_, 250 mM sucrose, 1% TritonX-100, 1% sodium deoxycholate, 100 μg/mL cycloheximide, 100 units/mL Superasin (Invitrogen), 0.25X protease inhibitor cocktail EDTA free (Roche, Sigma), and 1X phosphatase inhibitor cocktail (Santa Cruz Biotechnology). 9 to 10 OD Units measured at absorbance of 260 nm were loaded per linear sucrose gradient (20–47%). For genotype comparisons, the same OD units of lysate in same volume (700 μL) was layered per sucrose gradient (20 mM HEPES, pH 7.4, 200 mM KCl, 10 mM MgCl_2_, 100 µg/mL cycloheximide, RNase-free sucrose), and gradients were centrifuged at 35,000 RPM for 2 hours 45 min using a SW41Ti rotor (Beckman). Absorbance was followed at 254 nm. The relative area under the curve (AUC) was measured using the Peak Chart software (Brandel). Proteins were extracted from sucrose fractions using Trichloroacetic acid (TCA) precipitation. 105 μL of 3M TCA (Trichloroacetic acid) was added to each 750 μL fraction and vortexed on high for 20 to 30 sec and frozen at -80°C overnight. Samples were then thawed for 30 min followed by centrifugation at 20,000xg for 35 min at 4 °C and the supernatant was discarded. 1 mL of Acetone was added to each fraction and centrifuged at 20,000xg for 35 min at 4°C and the supernatant was discarded. The pellet was air dried and resuspended in 30 µL of 1X PBS and 30 µL of 2X Laemmli (SDS) buffer. For samples that were more acidic, pH was neutralized by adding 2 μL to 5 μL of 1M Tris and equal volume of 1M Tris was added to fractions that were being compared to keep the volumes consistent across samples. Samples were boiled for 8 min and same volume of each fraction was analyzed by western blotting.

### Immunohistochemistry

Paraffin embedded patient tissue sections were provided by the division of Pathology at Saint Louis University. Slides were warmed in a 60°C hybridization oven for 10 min. Slides were then deparaffinized by immersion in xylene for 5 min and the step was repeated. Slides were then dipped in graded alcohol sequentially from 100% (twice), 95% (twice) to 70% (twice) for 2 min each. Slides were washed by immersion in 1X PBS (three times) for 5 min each. Slides were quenched in 3% H_2_O_2_ in methanol for 10 min. Slides were then immersed in 1X PBS (three times) for 2 min each. Slides were immersed in a solution of 10 mM sodium citrate pH 6.0 for heat-induced antigen retrieval by boiling in a rice cooker for 30 min. Slides were then allowed to cool within that solution in a tightly closed container for 30 min. Slides were washed three times with 1X PBS for 2 min each. Slides were dried by wiping and a barrier was drawn around the tissue using a hydrophobic pen, (Fisher Scientific). Sections were handled carefully to avoid drying-out the tissue sections between each step. Blocking solution (1X PBS with 3% BSA and 0.1% Tween 20) was prepared right before use and vortexed well. Slides were placed in a humidified container, and then 300 μL of blocking solution was added to each section and incubated at RT for 30 min. Blocking solution was then removed and 200 μL of primary anti-eIF6 antibody (1:50, Santa Cruz Biotechnology) diluted in blocking solution was then added to each slide and incubated overnight at 4°C. The next day, slides were immersed in 1X PBS (three times) for 2 min each. The next set of steps were all conducted in the dark. Approximately 250 μL of secondary antibody-Alexa Fluor 546 (1:500, Invitrogen) diluted in blocking solution was then added to each section for 30 min at RT. Slides were washed three times in 1X PBS for 2 min each. Sections were mounted using Prolong Gold antifade mountant with DAPI stain (Invitrogen). Image J was used to quantitate fluorescence intensities for identical areas across all images. For each patient sample, two to four images were taken, and 3 distinct areas were measured per image. Scientist performing the analysis was blind to the tumor grade. **Ethics statement:** Patient tissue samples were provided by Dr. Zachary Hamilton (Division of Urology, Saint Louis University) as approved by IRB protocol# 32419.

### Ribosome Profiling

HCT116 cells of the indicated genotype were cultured to 60 to 70% cell density and were washed and scraped gently with ice cold PBS and snap frozen. Cells were then processed for Ribo-seq analysis at the RNA core at Case Western University. The appropriate volume of cell extract containing equal OD was treated with MNase, CaCl_2_ (5mM final concentration), and Turbo DNAse I and incubated at 25°C for 30 min; the digestion was stopped with SUPERase-inhibitor and placing on ice. The cell extract was then loaded onto a 15%-45% sucrose gradient to separate the polysomes by ultra-centrifugation at 41,000 RPM for 2 hours and 26 min at 4°C. The relevant fractions corresponding to the 80S monosome were collected and precipitated using ethanol and the RNA was isolated by phenol-chloroform extraction. In parallel, total RNA was isolated from each sample and subjected to the Turbo DNase I digestion and then RNA-seq libraries were constructed using NEB Next Ultra II directional RNA seq kit. The ribosome-protected fragments were subjected to denaturing polyacrylamide-mediated gel electrophoresis and were gel-extracted based on size 25nt to 35nt. The RNA was precipitated and subjected to the RiboMinus Ribosomal RNA depletion kit (Ambion). The rRNA-depleted RNA footprints were then subjected to de-phosphorylation. Adapter ligation, reverse transcription and subsequent cDNA amplification was performed using the NEXT-Flex smRNA-Seq Kit v3 (Perkin Elmer); cDNA library QC was performed by bioanalyzer before sequencing on an Illumina NovaSeq 6000 single-end for 100 cycles. Both Ribo-seq and RNA-seq reads were aligned to the human genome assembly GRCH38/hg38 using STAR (PMID: 23104886). RSEM for RNA quantification and RibosomeprofilingQC was used to quantify RPF orf (transcript level) count. Normalization of raw read counts for gene features was conducted by with DESeq2. TE calculation for WT and mutant using DESeq2 with generated adjusted p-values was performed using the equation: log_2_TE = log_2_(avg_count_RPF)-log_2_(avg_count_RNA). The appropriate software used for analysis of the plots and GO analysis are indicated in the legends. For analysis, we excluded genes that had 0 reads for one or more replicate samples.

## Notes

### Competing Interest Statement

The authors have declared no competing interest.

### Summary of Updates

New Abstract and revised Ribo-seq analysis.

## References

1 Panse, V. G. & Johnson, A. W. Maturation of eukaryotic ribosomes: acquisition of functionality. Trends Biochem Sci 35, 260–266 (2010). 10.1016/j.tibs.2010.01.001

2 Nerurkar, P. et al. Eukaryotic Ribosome Assembly and Nuclear Export. Int Rev Cell Mol Biol 319, 107–140 (2015). 10.1016/bs.ircmb.2015.07.002

3 Kater, L. et al. Visualizing the Assembly Pathway of Nucleolar Pre-60S Ribosomes. Cell 171, 1599–1610 e1514 (2017). 10.1016/j.cell.2017.11.039

4 Sulima, S. O., Hofman, I. J. F., De Keersmaecker, K. & Dinman, J. D. How Ribosomes Translate Cancer. Cancer Discov 7, 1069–1087 (2017). 10.1158/2159-8290.CD-17-0550

5 Jaako, P. et al. eIF6 rebinding dynamically couples ribosome maturation and translation. Nat Commun 13, 1562 (2022). 10.1038/s41467-022-29214-7

6 Brina, D., Grosso, S., Miluzio, A. & Biffo, S. Translational control by 80S formation and 60S availability: the central role of eIF6, a rate limiting factor in cell cycle progression and tumorigenesis. Cell Cycle 10, 3441–3446 (2011). 10.4161/cc.10.20.17796

7 Gandin, V. et al. Eukaryotic initiation factor 6 is rate-limiting in translation, growth and transformation. Nature 455, 684–688 (2008). 10.1038/nature07267

8 Miluzio, A., Beugnet, A., Volta, V. & Biffo, S. Eukaryotic initiation factor 6 mediates a continuum between 60S ribosome biogenesis and translation. EMBO Rep 10, 459–465 (2009). 10.1038/embor.2009.70

9 Miluzio, A. et al. Expression and activity of eIF6 trigger malignant pleural mesothelioma growth in vivo. Oncotarget 6, 37471–37485 (2015). 10.18632/oncotarget.5462

10 Basu, U., Si, K., Warner, J. R. & Maitra, U. The Saccharomyces cerevisiae TIF6 gene encoding translation initiation factor 6 is required for 60S ribosomal subunit biogenesis. Mol Cell Biol 21, 1453–1462 (2001). 10.1128/MCB.21.5.1453-1462.2001

11 Finch, A. J. et al. Uncoupling of GTP hydrolysis from eIF6 release on the ribosome causes Shwachman-Diamond syndrome. Genes Dev 25, 917–929 (2011). 10.1101/gad.623011

12 Weis, F. et al. Mechanism of eIF6 release from the nascent 60S ribosomal subunit. Nat Struct Mol Biol 22, 914–919 (2015). 10.1038/nsmb.3112

13 Gartmann, M., Blau, M., Armache, J. P., Mielke, T., Topf, M. & Beckmann, R. Mechanism of eIF6-mediated inhibition of ribosomal subunit joining. J Biol Chem 285, 14848–14851 (2010). 10.1074/jbc.C109.096057

14 Groft, C. M., Beckmann, R., Sali, A. & Burley, S. K. Crystal structures of ribosome anti-association factor IF6. Nat Struct Biol 7, 1156–1164 (2000). 10.1038/82017

15 Su, T. et al. Structure and function of Vms1 and Arb1 in RQC and mitochondrial proteome homeostasis. Nature 570, 538–542 (2019). 10.1038/s41586-019-1307-z

16 Ceci, M. et al. Release of eIF6 (p27BBP) from the 60S subunit allows 80S ribosome assembly. Nature 426, 579–584 (2003). 10.1038/nature02160

17 Brina, D. et al. eIF6 coordinates insulin sensitivity and lipid metabolism by coupling translation to transcription. Nat Commun 6, 8261 (2015). 10.1038/ncomms9261

18 Miluzio, A. et al. Impairment of cytoplasmic eIF6 activity restricts lymphomagenesis and tumor progression without affecting normal growth. Cancer Cell 19, 765–775 (2011). 10.1016/j.ccr.2011.04.018

19 Warren, A. J. Molecular basis of the human ribosomopathy Shwachman-Diamond syndrome. Adv Biol Regul 67, 109–127 (2018). 10.1016/j.jbior.2017.09.002

20 Wong, C. C., Traynor, D., Basse, N., Kay, R. R. & Warren, A. J. Defective ribosome assembly in Shwachman-Diamond syndrome. Blood 118, 4305–4312 (2011). 10.1182/blood-2011-06-353938

21 Grosso, S. et al. The pathogenesis of mesothelioma is driven by a dysregulated translatome. Nat Commun 12, 4920 (2021). 10.1038/s41467-021-25173-7

22 Shimamura, A. Molecular alterations governing predisposition to myelodysplastic syndromes: Insights from Shwachman-Diamond syndrome. Best Pract Res Clin Haematol 34, 101252 (2021). 10.1016/j.beha.2021.101252

23 Loreni, F., Mancino, M. & Biffo, S. Translation factors and ribosomal proteins control tumor onset and progression: how? Oncogene 33, 2145–2156 (2014). 10.1038/onc.2013.153

24 Challa, S. et al. Ribosome ADP-ribosylation inhibits translation and maintains proteostasis in cancers. Cell 184, 4531–4546 e4526 (2021). 10.1016/j.cell.2021.07.005

25 Menne, T. F. et al. The Shwachman-Bodian-Diamond syndrome protein mediates translational activation of ribosomes in yeast. Nat Genet 39, 486–495 (2007). 10.1038/ng1994

26 Tan, S. et al. Somatic genetic rescue of a germline ribosome assembly defect. Nat Commun 12, 5044 (2021). 10.1038/s41467-021-24999-5

27 Tan, S. et al. EFL1 mutations impair eIF6 release to cause Shwachman-Diamond syndrome. Blood 134, 277–290 (2019). 10.1182/blood.2018893404

28 Calamita, P. et al. SBDS-Deficient Cells Have an Altered Homeostatic Equilibrium due to Translational Inefficiency Which Explains their Reduced Fitness and Provides a Logical Framework for Intervention. PLoS Genet 13, e1006552 (2017). 10.1371/journal.pgen.1006552

29 Kennedy, A. L. et al. Distinct genetic pathways define pre-malignant versus compensatory clonal hematopoiesis in Shwachman-Diamond syndrome. Nat Commun 12, 1334 (2021). 10.1038/s41467-021-21588-4

30 Elliff, J. et al. Dynamic states of eIF6 and SDS variants modulate interactions with uL14 of the 60S ribosomal subunit. Nucleic Acids Res 51, 1803–1822 (2023). 10.1093/nar/gkac1266

31 Golob-Schwarzl, N. et al. Separation of low and high grade colon and rectum carcinoma by eukaryotic translation initiation factors 1, 5 and 6. Oncotarget 8, 101224–101243 (2017). 10.18632/oncotarget.20642

32 Luzha, J. et al. Expression of Eukaryotic Translation Initiation Factors in the Urothelial Carcinoma of the Bladder. Anticancer Res 43, 1437–1448 (2023). 10.21873/anticanres.16292

33 Lindstrom, M. S., Bartek, J. & Maya-Mendoza, A. p53 at the crossroad of DNA replication and ribosome biogenesis stress pathways. Cell Death Differ 29, 972–982 (2022). 10.1038/s41418-022-00999-w

34 Mah, L. J., El-Osta, A. & Karagiannis, T. C. gammaH2AX: a sensitive molecular marker of DNA damage and repair. Leukemia 24, 679–686 (2010). 10.1038/leu.2010.6

35 Ye, C. et al. BCCIP is required for nucleolar recruitment of eIF6 and 12S pre-rRNA production during 60S ribosome biogenesis. Nucleic Acids Res 48, 12817–12832 (2020). 10.1093/nar/gkaa1114

36 Hogan, B. L. & Korner, A. Ribosomal subunits of Landschutz ascites cells during changes in polysome distribution. Biochim Biophys Acta 169, 129–138 (1968). 10.1016/0005-2787(68)90014-2

37 Kaminskas, E. Serum-mediated stimulation of protein synthesis in Ehrlich ascites tumor cells. J Biol Chem 247, 5470–5476 (1972).

38 Frerichs, K. U. et al. Suppression of protein synthesis in brain during hibernation involves inhibition of protein initiation and elongation. Proc Natl Acad Sci U S A 95, 14511–14516 (1998). 10.1073/pnas.95.24.14511

39 Fan, H. & Penman, S. Regulation of protein synthesis in mammalian cells. II. Inhibition of protein synthesis at the level of initiation during mitosis. J Mol Biol 50, 655–670 (1970). 10.1016/0022-2836(70)90091-4

40 Wilker, E. W. et al. 14-3-3sigma controls mitotic translation to facilitate cytokinesis. Nature 446, 329–332 (2007). 10.1038/nature05584

41 Pyronnet, S., Pradayrol, L. & Sonenberg, N. A cell cycle-dependent internal ribosome entry site. Mol Cell 5, 607–616 (2000). 10.1016/s1097-2765(00)80240-3

42 Qin, X. & Sarnow, P. Preferential translation of internal ribosome entry site-containing mRNAs during the mitotic cycle in mammalian cells. J Biol Chem 279, 13721–13728 (2004). 10.1074/jbc.M312854200

43 Tanenbaum, M. E., Stern-Ginossar, N., Weissman, J. S. & Vale, R. D. Regulation of mRNA translation during mitosis. Elife 4 (2015). 10.7554/eLife.07957

44 Ingolia, N. T., Hussmann, J. A. & Weissman, J. S. Ribosome Profiling: Global Views of Translation. Cold Spring Harb Perspect Biol 11 (2019). 10.1101/cshperspect.a032698

45 Iwasaki, S. & Ingolia, N. T. The Growing Toolbox for Protein Synthesis Studies. Trends Biochem Sci 42, 612–624 (2017). 10.1016/j.tibs.2017.05.004

46 Ingolia, N. T., Ghaemmaghami, S., Newman, J. R. & Weissman, J. S. Genome-wide analysis in vivo of translation with nucleotide resolution using ribosome profiling. Science 324, 218–223 (2009). 10.1126/science.1168978

47 Guydosh, N. R. & Green, R. Dom34 rescues ribosomes in 3’ untranslated regions. Cell 156, 950–962 (2014). 10.1016/j.cell.2014.02.006

48 Young, D. J., Guydosh, N. R., Zhang, F., Hinnebusch, A. G. & Green, R. Rli1/ABCE1 Recycles Terminating Ribosomes and Controls Translation Reinitiation in 3’UTRs In Vivo. Cell 162, 872–884 (2015). 10.1016/j.cell.2015.07.041

49 Origanti, S., Cai, S. R., Munir, A. Z., White, L. S. & Piwnica-Worms, H. Synthetic lethality of Chk1 inhibition combined with p53 and/or p21 loss during a DNA damage response in normal and tumor cells. Oncogene 32, 577–588 (2013). 10.1038/onc.2012.84

50 Tan, T. Z. et al. Epithelial-mesenchymal transition spectrum quantification and its efficacy in deciphering survival and drug responses of cancer patients. Embo Mol Med 6, 1279–1293 (2014). 10.15252/emmm.201404208

51 Alamo, P. et al. Subcutaneous preconditioning increases invasion and metastatic dissemination in mouse colorectal cancer models. Dis Model Mech 7, 387–396 (2014). 10.1242/dmm.013995

52 Wang, H. et al. Epithelial-Mesenchymal Transition (EMT) Induced by TNF-α Requires AKT/GSK-3β-Mediated Stabilization of Snail in Colorectal Cancer. Plos One 8 (2013). ARTN e56664, 10.1371/journal.pone.0056664

53 Friedl, P., Locker, J., Sahai, E. & Segall, J. E. Classifying collective cancer cell invasion. Nat Cell Biol 14, 777–783 (2012). 10.1038/ncb2548

54 Pinzaglia, M. et al. EIF6 over-expression increases the motility and invasiveness of cancer cells by modulating the expression of a critical subset of membrane-bound proteins. BMC Cancer 15, 131 (2015). 10.1186/s12885-015-1106-3

55 Denny, S. K. et al. Nfib Promotes Metastasis through a Widespread Increase in Chromatin Accessibility. Cell 166, 328–342 (2016). 10.1016/j.cell.2016.05.052

56 Gao, G. Z. et al. The NFIB/CARM1 partnership is a driver in preclinical models of small cell lung cancer. Nature Communications 14 (2023). ARTN 363, 10.1038/s41467-023-35864-y

57 Barone, M. et al. Designed nanomolar small-molecule inhibitors of Ena/VASP EVH1 interaction impair invasion and extravasation of breast cancer cells. Proc Natl Acad Sci U S A 117, 29684–29690 (2020). 10.1073/pnas.2007213117

58 Carmona, G. et al. Lamellipodin promotes invasive 3D cancer cell migration via regulated interactions with Ena/VASP and SCAR/WAVE. Oncogene 35, 5155–5169 (2016). 10.1038/onc.2016.47

59 Damiano-Guercio, J. et al. Loss of Ena/VASP interferes with lamellipodium architecture, motility and integrin-dependent adhesion. Elife 9 (2020). ARTN e55351, 10.7554/eLife.55351

60 Sanvito, F. et al. Expression of a highly conserved protein, p27BBP, during the progression of human colorectal cancer. Cancer Res 60, 510–516 (2000).

61 Gatza, M. L., Silva, G. O., Parker, J. S., Fan, C. & Perou, C. M. An integrated genomics approach identifies drivers of proliferation in luminal-subtype human breast cancer. Nat Genet 46, 1051–1059 (2014). 10.1038/ng.3073

62 Smith, P. R., Pandit, S. C., Loerch, S. & Campbell, Z. T. The space between notes: emerging roles for translationally silent ribosomes. Trends in Biochemical Sciences 47, 477–491 (2022). 10.1016/j.tibs.2022.02.003

63 Brown, A., Baird, M. R., Yip, M. C. J., Murray, J. & Shao, S. Structures of translational inactive mammalian ribosomes. Elife 7 (2018). ARTN e40486, 10.7554/eLife.40486

64 Wells, J. N. et al. Structure and function of yeast Lso2 and human CCDC124 bound to hibernating ribosomes. Plos Biol 18 (2020). ARTN e3000780,10.1371/journal.pbio.3000780

65 Van Dyke, N., Baby, J. & Van Dyke, M. W. Stm1p, a ribosome-associated protein, is important for protein synthesis under nutritional stress conditions. Journal of Molecular Biology 358, 1023–1031 (2006). 10.1016/j.jmb.2006.03.018

66 Wang, Y. N. J. et al. Lso2 is a conserved ribosome-bound protein required for translational recovery in yeast. Plos Biol 16 (2018). ARTN e2005903, 10.1371/journal.pbio.2005903

67 Pisareva, V. P., Skabkin, M. A., Hellen, C. U. T., Pestova, T. V. & Pisarev, A. V. Dissociation by Pelota, Hbs1 and ABCE1 of mammalian vacant 80S ribosomes and stalled elongation complexes. Embo Journal 30, 1804–1817 (2011). 10.1038/emboj.2011.93

68 van den Elzen, A. M., Schuller, A., Green, R. & Seraphin, B. Dom34-Hbs1 mediated dissociation of inactive 80S ribosomes promotes restart of translation after stress. EMBO J 33, 265–276 (2014). 10.1002/embj.201386123

69 Adham, I. M. et al. Disruption of the pelota gene causes early embryonic lethality and defects in cell cycle progression. Mol Cell Biol 23, 1470–1476 (2003). 10.1128/MCB.23.4.1470-1476.2003

70 Jungers, C. F., Elliff, J. M., Masson-Meyers, D. S., Phiel, C. J. & Origanti, S. Regulation of eukaryotic translation initiation factor 6 dynamics through multisite phosphorylation by GSK3. J Biol Chem (2020). 10.1074/jbc.RA120.013324

71 Wyler, E., Wandrey, F., Badertscher, L., Montellese, C., Alper, D. & Kutay, U. The beta-isoform of the BRCA2 and CDKN1A(p21)-interacting protein (BCCIP) stabilizes nuclear RPL23/uL14. FEBS Lett 588, 3685–3691 (2014). 10.1016/j.febslet.2014.08.013

72 Ting, Y. H. et al. Bcp1 Is the Nuclear Chaperone of Rpl23 in Saccharomyces cerevisiae. J Biol Chem 292, 585–596 (2017). 10.1074/jbc.M116.747634

73 Roshan, P. et al. An Aurora B-RPA signaling axis secures chromosome segregation fidelity. Nat Commun 14, 3008 (2023). 10.1038/s41467-023-38711-2

