## Supplementary material for "Modulation of ribosomal subunit associations by eIF6 is critical for mitotic exit and cancer progression": All Suppl. images

**Figure S1**

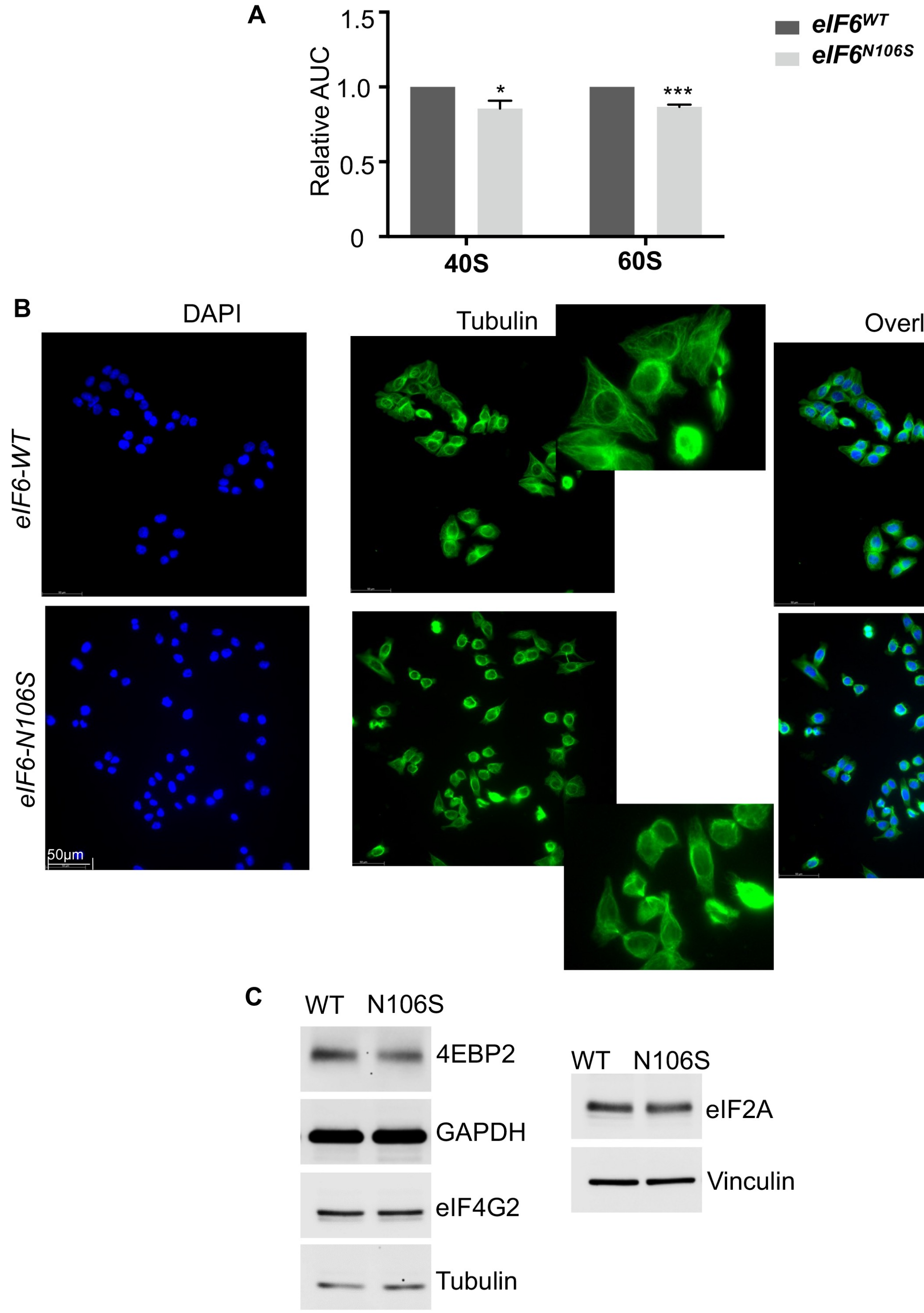


**Figure S1. Minimal decrease in 60S and 40S levels and mild changes in cell morphology in eIF6-N106S mutant.**

A) Plot shows the relative area under the curve (AUC) measurements for 40S and 60S ribosomal peaks in eIF6-WT and eIF6-N106S mutant. Plot shows standard error of the mean from four independent experiments with a significant difference for 40S ($p$=0.032) and 60S ($p$=0.00012) as determined using an unpaired two-tailed $t$ test. B) Representative immunofluorescence (IF) images of fixed asynchronous cells stained with DAPI (nuclear marker) or treated with anti-α-Tubulin antibody. Enlarged inset shows the lack of spread in Tubulin network and altered cell to cell contact in mutant compared to WT. Images represent three independent experiments. C) Representative western blots were probed with the indicated antibodies. Blots were also probed with anti-Vinculin, anti-β-Tubulin or anti-GAPDH antibodies as loading controls. Data are representative of three independent experiments.

### Figure S2

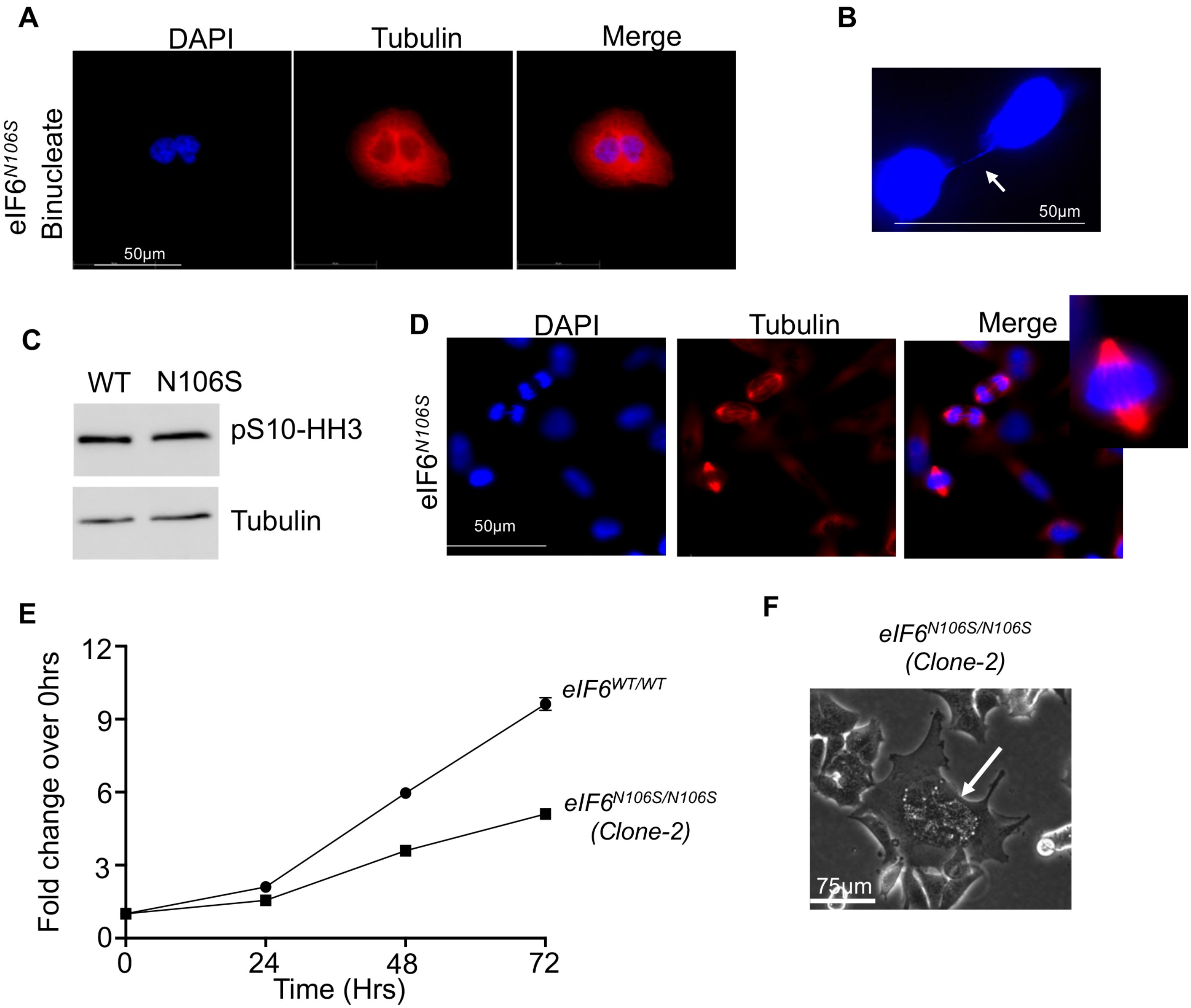


**Figure S2. Spindle bipolarity and mitotic entry unaltered in the eIF6-N106S mutant and delayed cell proliferation and mitotic catastrophe observed in the second clone of eIF6-N106S mutant.** A) Representative immunofluorescence (IF) images show increased presence of binucleated cells for the eIF6-N106S mutant. B) IF image shows daughter nuclei (stained with DAPI) tethered by chromatin bridges (white arrow). C) Western blot represents serine-10 phosphorylated Histone H3 levels in mitotic cells synchronized by nocodazole. Blots were also probed for Tubulin as loading control. D) Representative immunofluorescence (IF) images show bipolar spindles in mitotic cells of asynchronous eIF6-N106S mutant cell population. E) Plot indicates the fold change in cell proliferation rates relative to 0 hours of plating as measured by the MTS assay for a second clone of eIF6-N106S in comparison to eIF6-WT. Plot indicates standard error of the mean from three independent experiments with triplicate wells measured per experiment. The difference between eIF6-WT and eIF6-N106S was significant at 24 hours ($p$=0.00000002), 48 hours ($p$=0. 00000063) and 72 hours ($p$=0.00000002) of growth as determined by an unpaired two-tailed $t$ test. F) Brightfield image shows the presence of mitotic catastrophe in a multinucleated cell with vacuoles (white arrow) in the second clone of eIF6-N106S mutant.

**Figure S3**

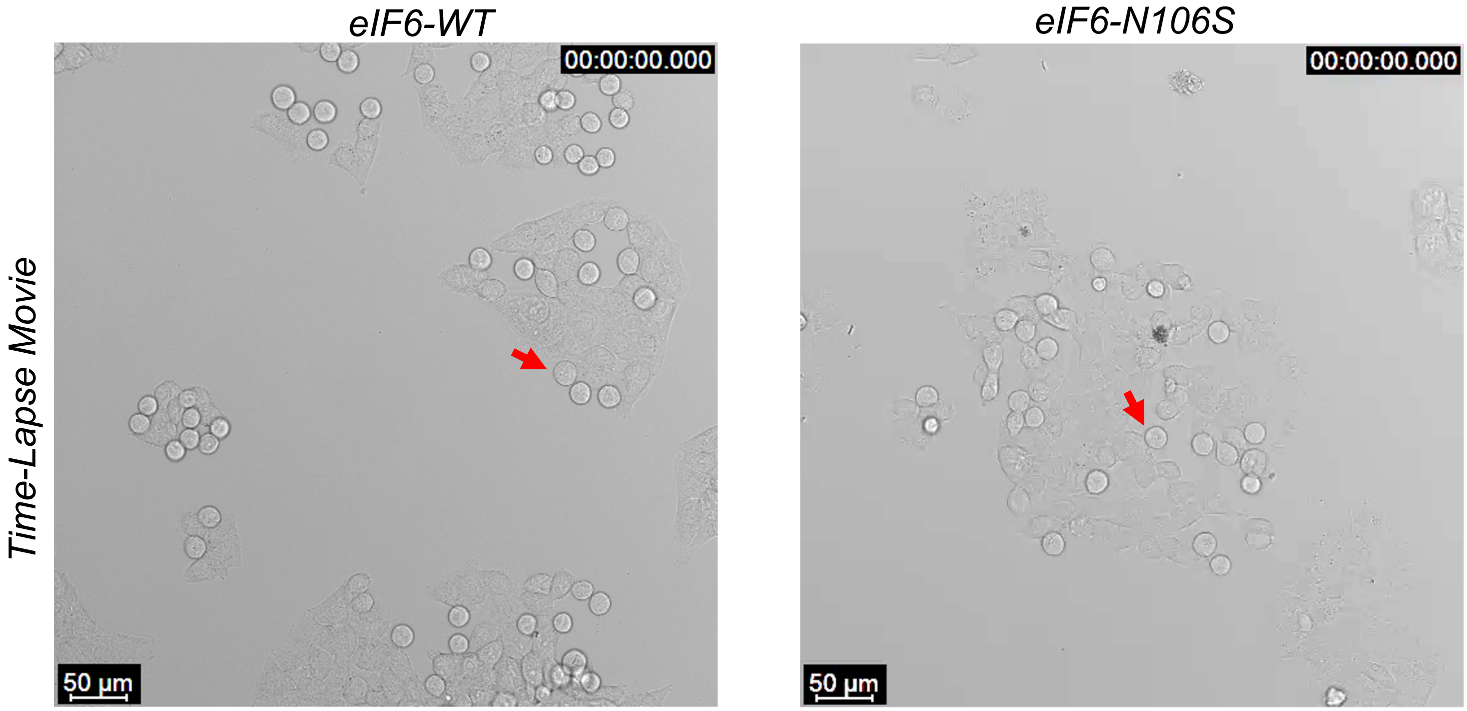


**Figure S3. Delayed mitotic exit in eIF6-N106S mutant.** A) Representative brightfield time-lapse confocal videos show the timing of cell cycle progression from metaphase to G1. Red arrows indicate the cells that were highlighted in the main figure.

#### Figure S4

**A**

| Sample- RNA | Total_reads | Average_read_length | Unique_map_rate | Multiple_map_rate |
|---|---|---|---|---|
| S01 (WT-1) | 90443822 | 76 | 33.42% | 66.06% |
| S02 (WT-2) | 72100419 | 76 | 38.34% | 60.61% |
| S03 (WT-3) | 67404104 | 76 | 34.80% | 62.58% |
| S04 (Mutant-1) | 83352910 | 76 | 35.23% | 63.74% |
| S05 (Mutant-2) | 73685702 | 76 | 38.69% | 60.12% |
| S06 (Mutant-3) | 57326742 | 76 | 35.39% | 62.23% |

**B**

| Sample-RPF | Total_reads | Average_read_length | Unique_map_rate | Multiple_map_rate |
|---|---|---|---|---|
| SMT1 (Mutant-1) | 6564802 | 32 | 61.76% | 36.39% |
| SMT2 (Mutant-2) | 7595767 | 32 | 64.97% | 33.33% |
| SMT3 (Mutant-3 | 7918603 | 31 | 64.07% | 34.06% |
| SW1 (WT-1) | 4685932 | 32 | 60.26% | 37.96% |
| SW3 (WT-3) | 7074993 | 35 | 63.88% | 34.73% |

**Figure S4. Quality control of the RNA-Seq and Ribo-Seq data.** A) Table shows the total reads, average read length, unique mapped read rates and multiple mapped reads rate for RNA-seq library of three independent WT and three independent eIF6-N106S mutant samples. B) Table shows the total reads, average read length, unique mapped reads rate and multiple mapped reads rate for RPF-seq library of two independent WT and three independent eIF6-N106S mutant samples. One of the WT-samples (SW2) did not pass quality control for RPF analysis and was not used for analysis.

#### Figure S5

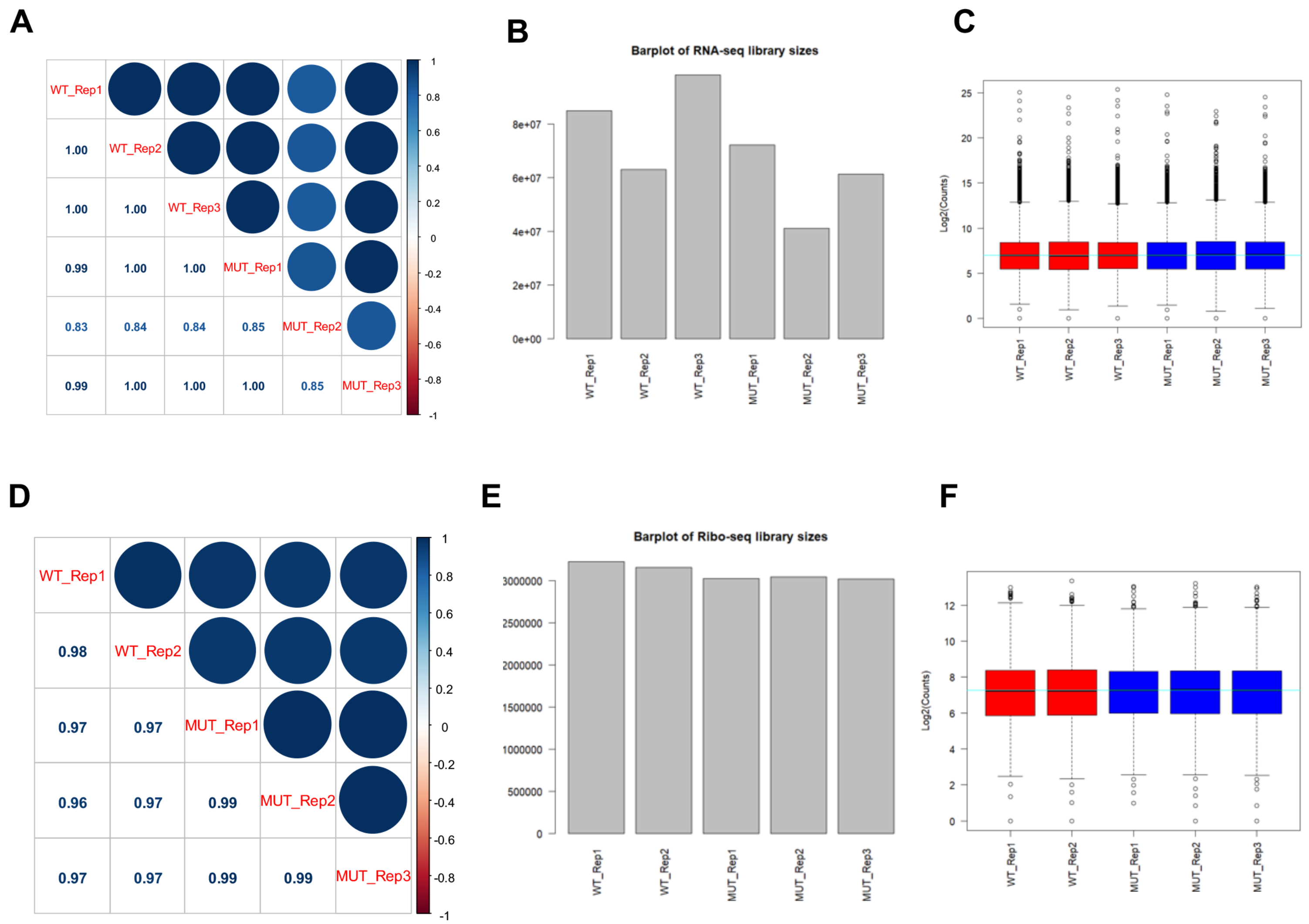


**Figure S5. Quality control of the RNA-Seq and Ribo-Seq data.** **A**) Correlation analysis of RNA-Seq results for biological replicates. Correlation matrix showing Pearson correlation coefficients calculated for pair-wise comparisons of numbers of raw counts of RNA-Seq reads for all expressed genes among 5 RNA-Seq libraries. B) A bar graph showing library size (normalized read count) for each RNA-Seq library. C) A boxplot showing the distribution of gene read count in each each RNA-Seq library. D) Correlation analysis of Ribo-Seq (RPF) results for biological replicates. Correlation matrix showing Pearson correlation coefficients calculated for pair-wise comparisons of numbers of raw counts of RPF reads for all expressed genes among 5 RPF libraries. E) A bar graph showing library size (normalized read count) for each Ribo-Seq (RPF) library. F) A boxplot showing the distribution of gene read count in each Ribo-Seq (RPF) library.

### Figure S6

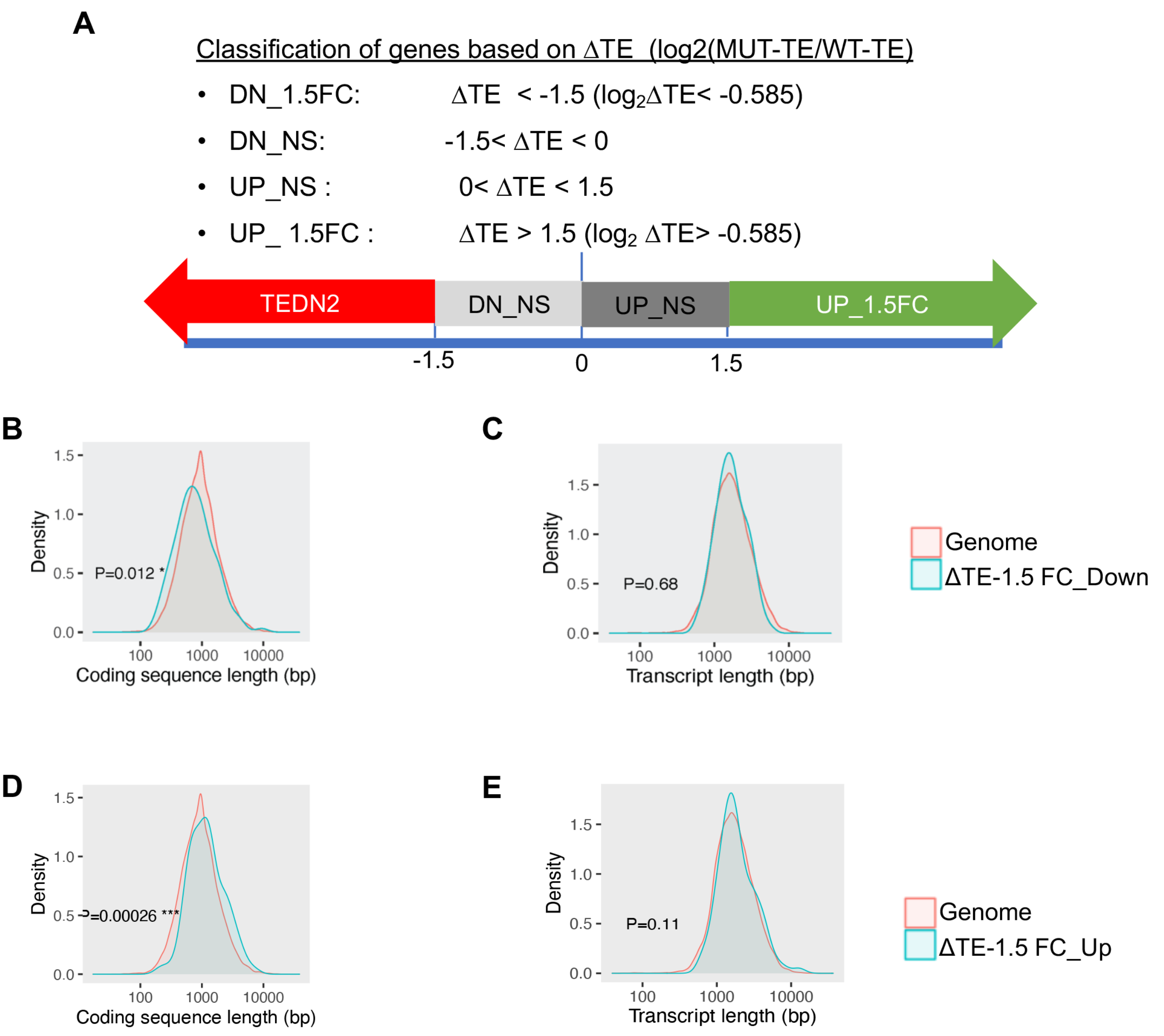


**Figure S6. Ribo-Seq data analyses.** A) Graphic shows the equation for the calculation of ΔTE and depicts the classification of genes based on ΔTE. B and C) Density plots show the density of transcripts that are significantly downregulated by 1.5-fold or more in mutant compared to WT and relative to the genome. Comparisons of density of coding region length (B) and total transcript length (inclusive of 5'UTR, coding and 3'UTR) (C) are shown. D and E). Density plots show the density of transcripts that are significantly upregulated by 1.5-fold or more in mutant compared to WT and relative to the genome. Comparisons of density of coding region length (D) and total transcript length (inclusive of 5'UTR, coding and 3'UTR) (E) are shown.

### Figure S7

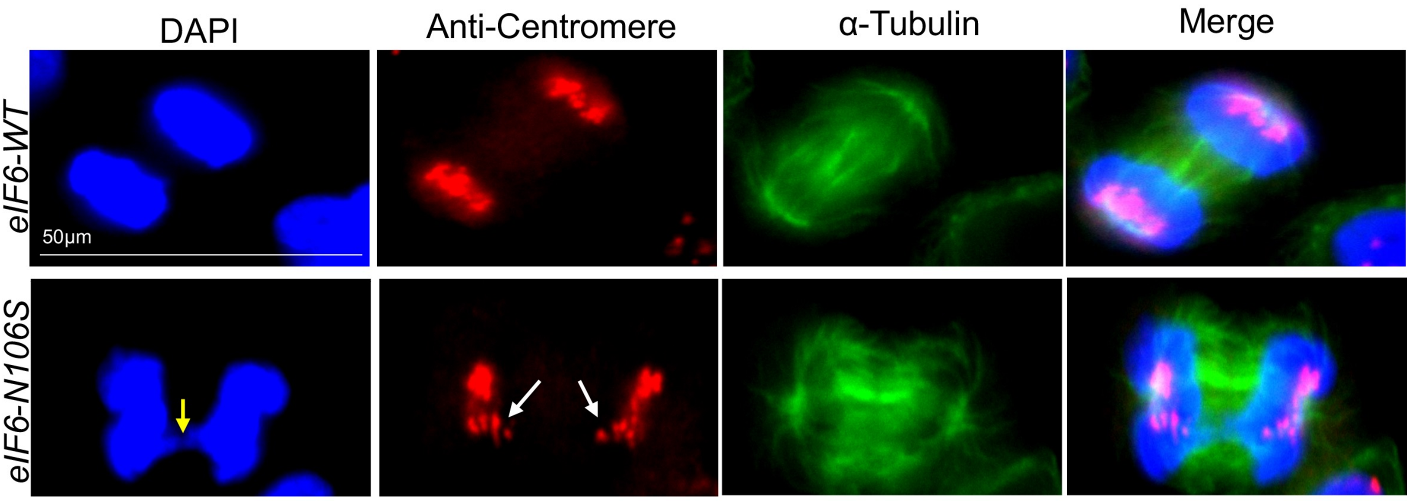


**Figure S7. Chromosome misalignment in the eIF6-N106S mutant.** A) Representative immunofluorescence (IF) images show increased presence of cells in late mitosis. Cells were stained with DAPI (blue) or anti-centromere (CREST) (red) or α-Tubulin (green) antibodies. Yellow arrow indicates the anaphase bridge and white arrows indicate the misaligned chromosomes as shown by anti-centromere antibody staining. Data represents three independent experiments.

#### Figure S8

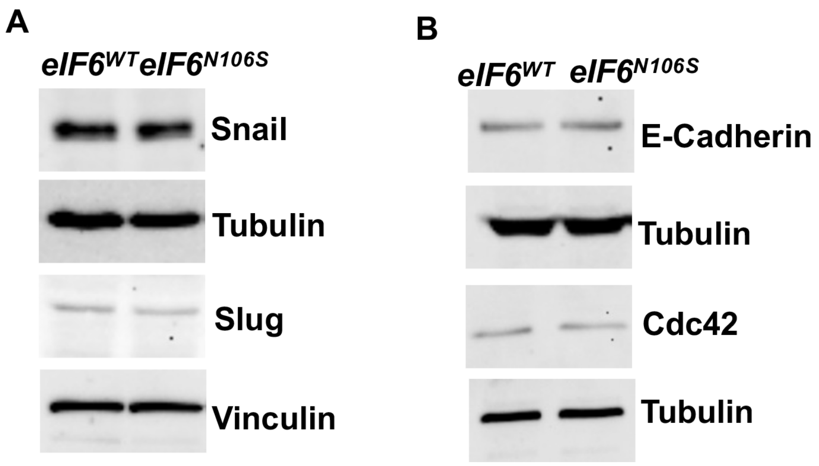


**Figure S8. Analysis of EMT markers.** A and B) Representative western blots probed with the indicated antibodies are shown. Blots were also probed with anti-Tubulin or anti-Vinculin antibodies as loading controls. Data represents three independent experiments.

**Figure S9**

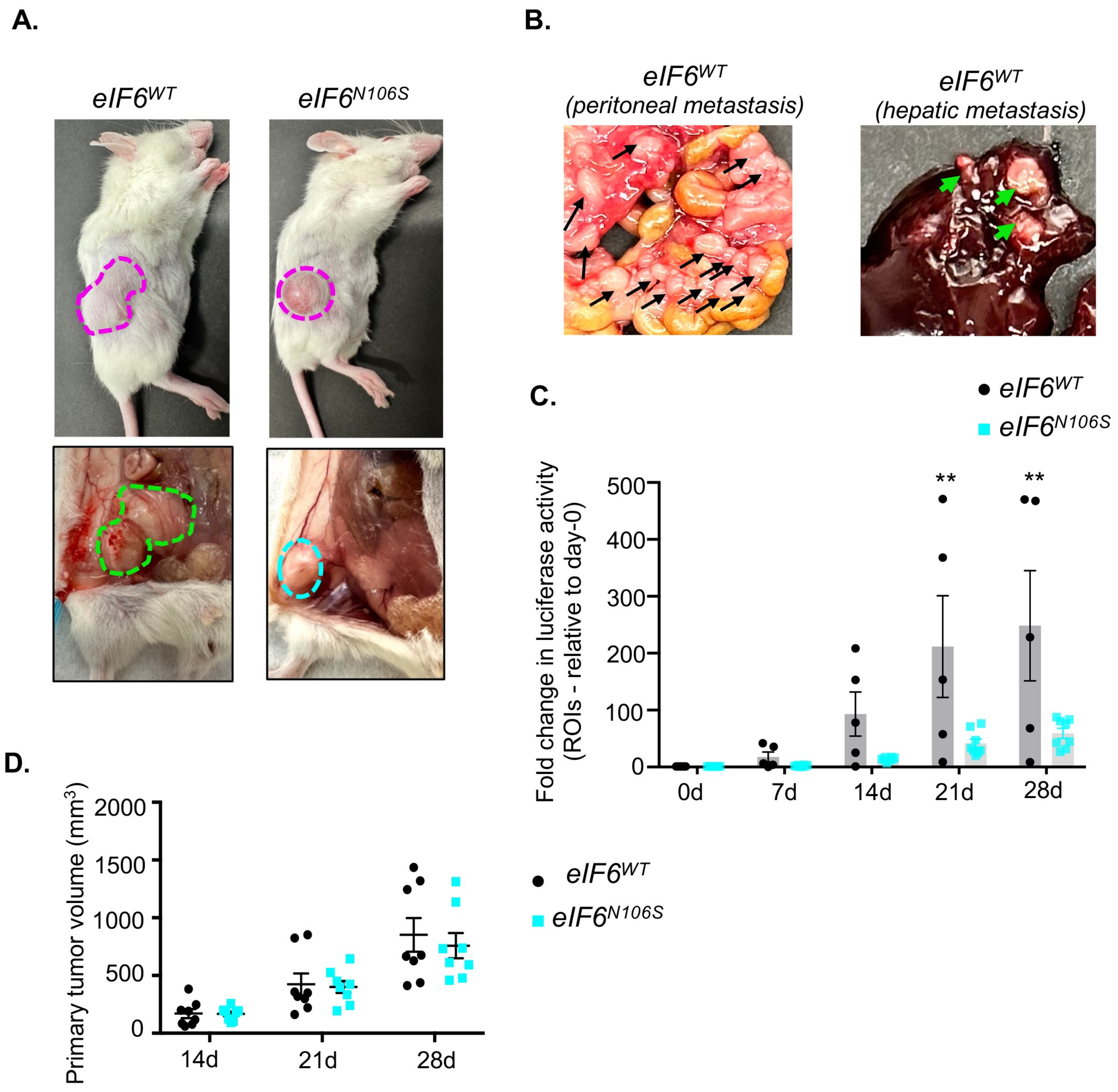


**Figure S9. eIF6-N106S mutant tumors do not exhibit invasion *in vivo*.** A) Images represent the spread-out growth (pink and green dotted lines) of subcutaneous tumors of eIF6-WT mice relative to the localized growth of eIF6-N106S cells (pink and blue dotted lines) on day 28. Images on top represent the external view of tumor and bottom represent the internal view. B) Tissue images show the presence of several peritoneal metastatic regions (black arrows) in the intestines and hepatic metastases (green arrows) in mice carrying eIF6-WT tumors on day 28. C) Plot shows the fold change in luciferase activity at Days 7, 14, 21 and 28 relative to Day 0 for n=8 eIF6-WT and n=8 eIF6-N106S tumors. Significant differences were determined using two-way ANOVA using Sidak's multiple comparisons test with $p$=0.0038 at Day 21 and $p$=0.0011 at Day 28. D) Plot represents primary tumor volumes measured for n=8 eIF6-WT and n=8 eIF6-N106S tumors using calipers. However, the measurements are skewed by the irregular shape of some of the WT tumors and part of tumor growing inwards.

#### Figure S10

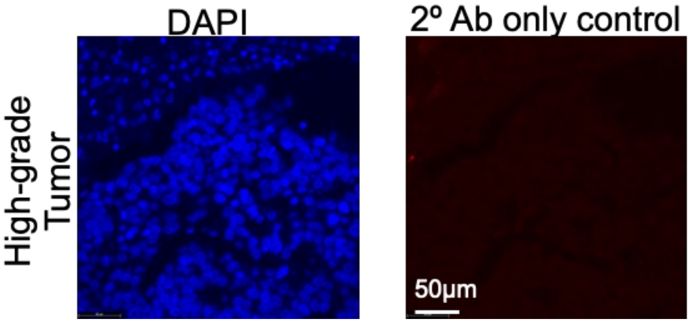


**Figure S10.** Representative IF image of patient-derived high-grade tumor section probed with DAPI (nuclear stain) or secondary antibody control only. Data represents three independent experiments.
